# Foamy microglia link oxylipins to disease progression in multiple sclerosis

**DOI:** 10.1101/2024.10.18.619040

**Authors:** Daan van der Vliet, Xinyu Di, Tatiana M. Shamorkina, Anto Pavlovic, Iris A.C.M. van der Vliet, Yingyu Zeng, Will Macnair, Noëlle van Egmond, J.Q. Alida Chen, Aletta M.R. van den Bosch, Hendrik J. Engelenburg, Matthew R.J. Mason, Claire Coulon-Bainier, Berend Gagestein, Elise Dusseldorp, Marco van Eijk, Uwe Grether, The Netherlands Brain Bank, Amy C. Harms, Thomas Hankemeier, Ludovic Collin, Albert J.R. Heck, Inge Huitinga, Mario van der Stelt

## Abstract

Multiple sclerosis (MS) is a neuroinflammatory disease characterized by expanding demyelinating lesions, leading to severe and irreversible disability. The mechanisms driving lesion expansion, however, remain poorly understood. Here, using a multi-omics approach, we identified foamy microglia as primary contributors to the molecular profile of lesions and disease progression in secondary progressive MS. Lesions with foamy microglia are marked by the accumulation of cholesterol esters, bismonoacylglycerolphosphates (BMP), and oxylipins, along with high B-cell infiltration, increased levels of immunoglobulin G1, and elevated expression of Fcγ- and complement receptors. Lesions with foamy GPNMB^+^-microglia display markers of enhanced phagocytosis, lipid metabolism, lysosomal dysfunction, and antigen presentation, but lack classical pro-inflammatory markers. Our data suggest that sustained phagocytosis of myelin overwhelms microglial endo-lysosomal capacity, leading to lipid droplet and oxylipin formation. This microglial phenotype may induce further recruitment of adaptive immune cells, axonal damage, drive lesion expansion and prevent remyelination. Monoacylglycerol lipase, involved in producing oxylipin precursors, was identified as a potential therapeutic target to disrupt this cycle and prevent chronic lesion expansion.

## Introduction

Multiple sclerosis (MS) is a complex and debilitating neurological disorder characterized by the formation of inflammatory demyelinating lesions throughout the central nervous system. Unresolved lesions are associated with extensive axonal damage, gliosis, and cell death that result in severe, often irreversible, disability in patients.^1,2^ Despite significant advances in the development of disease-modifying therapies that are effective in the early phases of MS, there remains a critical unmet need for treatments that can halt disease progression^3^.

The progression of MS is notably heterogeneous, both in clinical presentation and in the underlying pathology observed in post-mortem studies^4^. This heterogeneity is reflected in the diverse pathology of MS lesions, which vary in size, cellular composition, immune cell activity, and the extent of demyelination.^5–7^ These observations have led to the classification of MS lesions into four distinct subtypes: active, mixed active/inactive, inactive, and remyelinated lesions (Figure 1).^1^ Active and mixed active/inactive lesions (AL and ML, respectively), marked by active microglial cells expressing high levels of HLA-DR/DQ/DP, suggesting ongoing demyelination, whereas inactive lesions (IL) are characterized by severe astrogliosis and a lack of remyelination, indicating a failure of repair processes and permanent axonal damage. Remyelinated lesions (RL), often referred to as “shadow plaques”, exhibit partial repair, which is thought to contribute to the relapsing-remitting nature of early-stage MS. However, capacity for remyelination diminishes with age, often failing in the progressive stages of the disease, leading to chronic loss of neurological function.^3,8–10^ However, the molecular mechanisms determining why some lesions resolve while other chronically expand remain elusive.

**Fig. 1.**
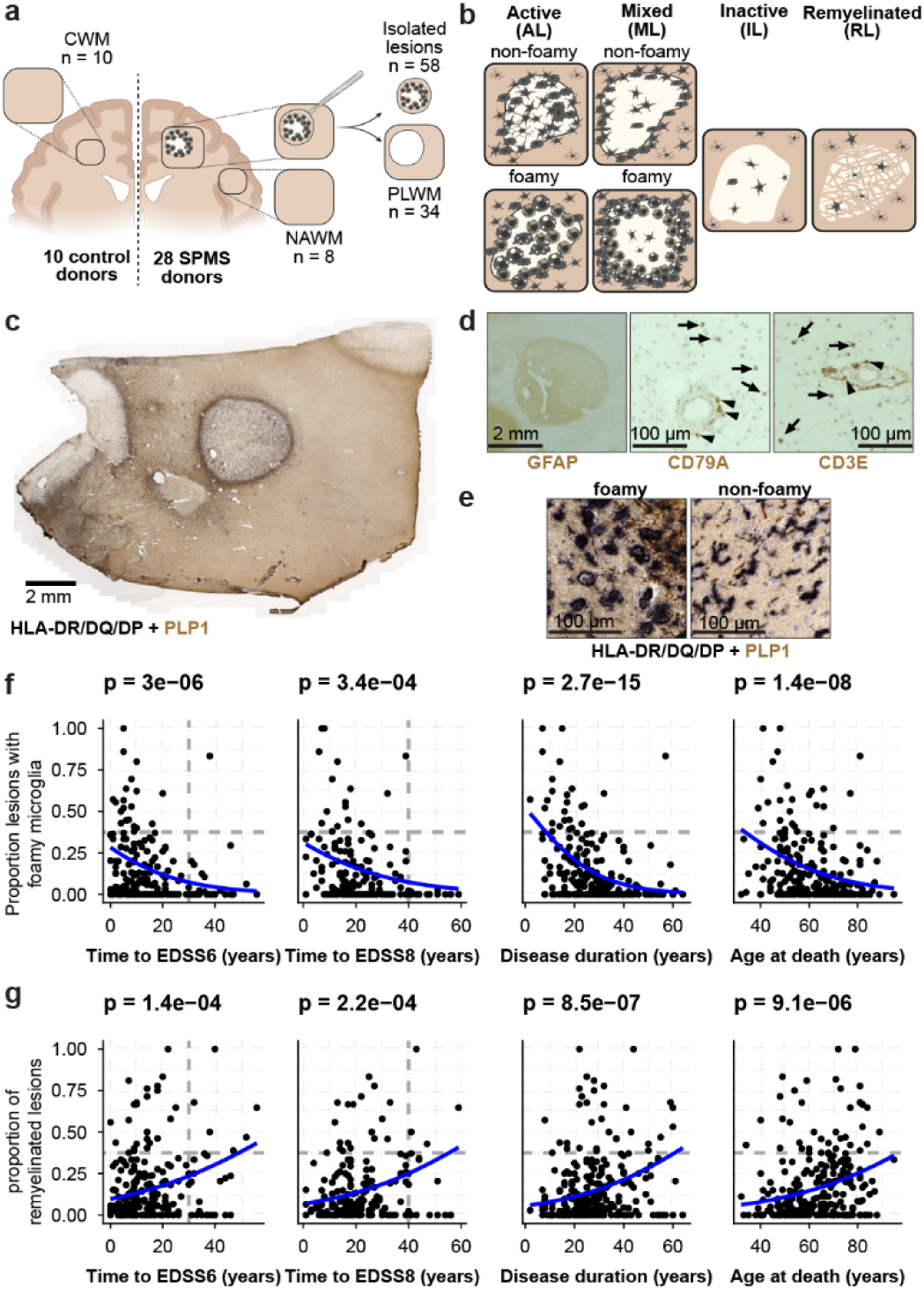
Foamy microglia associate with faster disease progression. **a,** Tissue sampling in this study. **b,** Graphical representation of white matter lesions. ALs and MLs are subcategorized based on the foamy phenotype. **c-d,** Histological analysis of a representative white matter lesion, stained for PLP1 (myelin) and HLA-DR/DQ/DP (microglia), GFAP (astrocytes), CD79A (B-cells) or CD3 (T-cells) (**d**). Arrows indicate parenchymal cells, while arrowheads indicate perivascular cells. **e,** Representative images of foamy (left) or non-foamy microglia (right). **f-g,** Proportion of lesions with foamy microglia (**f**) or remyelinated lesions (**g**) associates with disease course (N=250 MS patients). Associations were estimated using quasibinomial generalized linear modelling with a likelihood ratio test and Bonferroni multiple testing correction.

A particularly intriguing aspect of MS pathology is the presence of foamy microglia within ALs and MLs. These microglia, characterized by their lipid-laden appearance^6,11–15^, are associated with increased axonal damage, higher levels of neurofilament light chain (NFL) in the cerebrospinal fluid (CSF), and greater overall lesion load, indicating a detrimental effect of foamy microglia in MS lesions.^5,6,16^ However, foamy microglia have also been described as anti-inflammatory or immunosuppressed.^11,15^ The exact role of foamy microglia in lesion etiology is currently still unclear.

Here, we have addressed the these gaps, by conducting a comprehensive multi-omics analysis, integrating lipidomics, transcriptomics and proteomics, to systematically profile well-characterized post-mortem white matter lesions from MS patients. Our findings highlight the significant role of foamy microglia in determining the molecular profile of MS lesions. Lesion with foamy microglia were characterized by cholesterol-ester and BMP accumulation, and the production of oxidized free fatty acids (oxylipins). In contrast, lesions with ramified microglia are more closely associated with successful remyelination. We identified possible drug targets to address the foamy phenotype using activity-based protein profiling. Lastly, we show that oxylipins in cerebrospinal fluid associate with the foamy phenotype. Our findings suggest that modulating lipid metabolism in foamy microglia may offer a promising therapeutic strategy to halt disease progression in MS.

## Main

The heterogeneity of MS disease progression is reflected by diverse post-mortem white matter (WM) pathology^6,7^. Thus, by studying the molecular profile of diverse WM lesions across a diverse MS cohort we hypothesized to uncover molecular patterns related to disease progression. To this end, we isolated fresh-frozen subcortical white matter samples from 28 individuals diagnosed with secondary progressive MS with diverse white matter pathology (Figure S1c) and 10 controls from the Netherlands Brain Bank (NBB) (Figure 1a-e). A total of 110 samples were isolated, covering 58 active, mixed active/inactive, inactive and remyelinated lesions (AL, ML, IL, and RL, respectively), alongside 52 samples from normal appearing white matter (NAWM), peri-lesional white matter (PLWM), and control white matter (CWM). We further subcategorized ALs and MLs based on microglial morphology: either foamy or non-foamy (Figure 1e)^6,11,12,14^. To assess the clinical relevance of this subcategorization, we coupled pathological data from all MS patients in the NBB (N=250) to their disease trajectory. The proportion of ALs and MLs with foamy microglia in the donor’s white matter was associated with faster progression (Figure 1f, S1d-e). In contrast, ALs and MLs with non-foamy microglia did not correlate with progression, while RLs correlated with slower progression (Figure 1g, S1d,e). These associations were also observed in this study’s cohort (N=28, Figure S1f-i).

While lipids are central to the biology of myelin^17^ and foamy microglia^15^, this class of biomolecules has been understudied compared to gene expression^18–24^ in the context of MS. Therefore, we set out to perform a comprehensive targeted lipidomics analysis of MS lesions. We measured 712 lipid species across 31 lipid classes using targeted LC-MS/MS. The analysis revealed that all MS lesions had a distinct lipid composition compared to CWM, NAWM and PLWM, with variability driven by lesion presence/absence and microglial morphology (Figure 2a-d, S2).

**Fig. 2.**
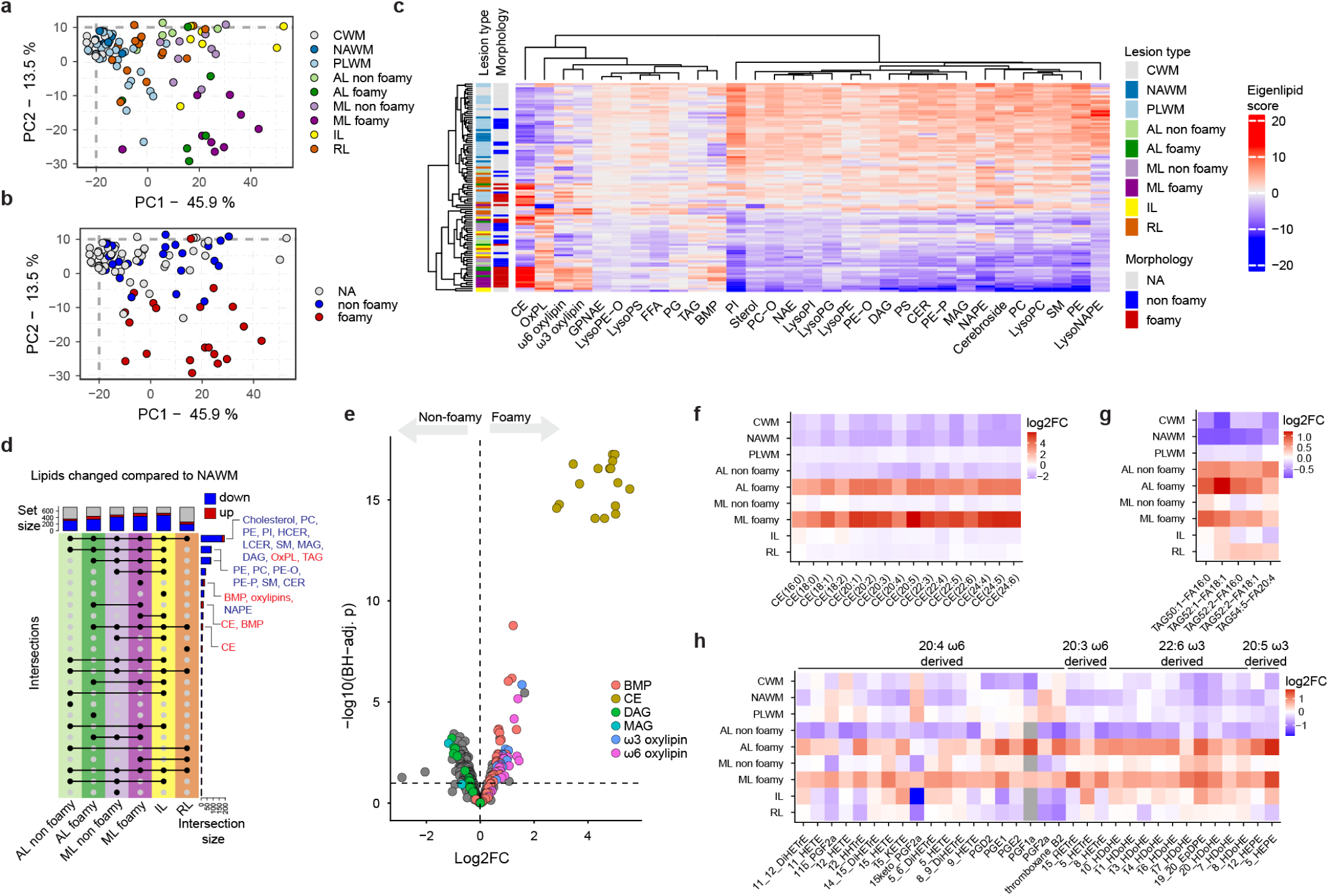
Lipidomics reveals general loss of lipids in lesions and a distinct lipid-profile of foamy microglia. **a-b,** Principal component analysis (PCA) of the lipidomics dataset coloured by (**a**) lesion type or (**b**) morphology. NA indicates samples without microglial activation. **c,** Clustered heatmap depicting the first principal component for each lipid class (the “eigenlipid”). Hierarchical clustering was performed using euclidean distances. **d,** Upset plot depicting the overlap of differentially abundant lipids for each lesion type compared to NAWM. **e,** Differentially abundant lipids between lesions with foamy-compared to lesions with non-foamy microglia. p-values are calculated using limma with BH-correction for multiple testing (FDR<10%). **f-h,** Normalized mean abundance of lipid classes CE (**f**), TAG (**g**) or oxylipins (**h**) across lesion types.

Myelin-associated lipids, such as cholesterol, phospholipids and sphingolipids, were significantly reduced in all lesion types (Figure 2c-d, S2a-e). These changes were mainly associated with the extent of demyelination, as the decrease of these lipids was much less in remyelinated lesions. The ratio between cholesterol and its derivatives increased (Figure S2f), indicating a shift from cholesterol synthesis to cholesterol recycling and oxidation in lesions^25^. Oxidized phospholipids (oxPLs), particularly those derived from poly-unsaturated acyl chains, were elevated in all lesion types, but were unlikely to directly induce demyelination at the observed levels (Figure S3a-e) as previously suggested^26,27^.

Lesions with foamy microglia exhibited increased cholesterol esters (CE) and triacylglycerides (TAG), consistent with their lipid-laden phenotype (Figure 2f-g)^11,13^. Notably, bismonoacylglycerolphosphates (BMPs, Figure S3f) and oxylipins (Figure 2h) were also strongly increased in the foamy lesions, which suggested an enhanced lysosomal processing and inflammatory state, respectively^28–30^.

To directly assess the inflammatory status of the lesions, we performed a cytokine screening of ML with either foamy or non-foamy microglia compared to NAWM and CWM. Notably, prominent inflammatory markers, like TNFα, IL6, IFNγ and IL1β, were not different between non-foamy and foamy lesions (Figure S4). Only three cytokines, associated with apoptosis (FasL)^31^, adaptive immune system signaling^32^ (CD8+ T-cells, CCL5) and cellular senescence (SerpinE1)^33^ were significantly upregulated in foamy lesions compared to non-foamy.

To better characterize the biological state of different white matter MS lesions, we performed bulk RNA sequencing and proteomics to capture both gene expression and post-transcriptional processes (Figure 3). We obtained expression data for 16,651 genes and 3,237 proteins, which generally showed poor correlation (Figure S5a-c). Notably, structural axonal proteins, whose gene transcripts primarily reside in neuronal somas, did not correlate well with protein levels, while proteins expressed by oligodendrocytes and microglia showed stronger correlation with their gene transcripts (Figure S5a-c). The primary drivers of variance in the transcriptomics and proteomics datasets were lesion presence/absence and the foamy phenotype (Figure S6), alike the lipidomics data. Most differentially expressed genes and proteins were consistent across all lesion types (Figure 3a,c), reflecting general lesion features such as demyelination (Figure S7), oligodendrocyte loss and astrogliosis. In addition, lesions with foamy microglia had a distinct transcriptome (Figure 3b) and proteome (Figure 3d) from those with non-foamy microglia. We observed increased expression of genes and proteins involved in cholesterol recycling, lipid droplet formation, oxylipin production and ferroptosis (Figure S8, S9), aligning well with the lipidomics dataset.

**Fig. 3.**
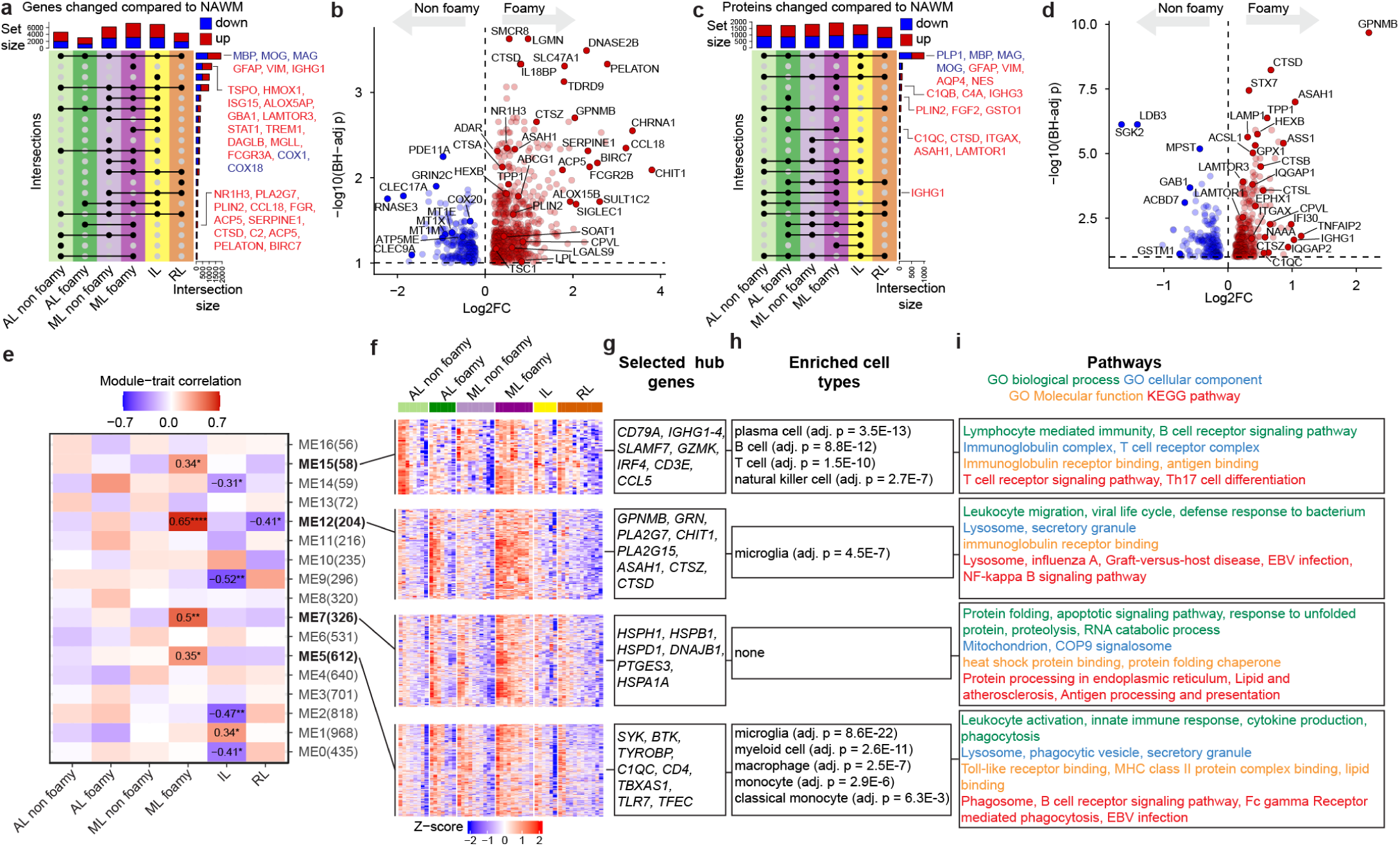
Lesions with foamy microglia have disturbed lipid metabolism and immune regulation. **a+c,** Upset plots depicting the overlap of differentially expressed genes (DEGs) (**a**) or differentially abundant proteins (DAPs) (**c**) for each lesion type compared to NAWM. **b+d,** DEGs (**b**) or DAPs (**d**) between lesions with foamy compared to lesions with non-foamy microglia. p-values were calculated using limma with BH-correction for multiple testing (FDR<10%). **e-j,** Weighted gene co-expression network analysis (WGCNA) divided the transcriptome of lesions into 17 modules of co-expressed genes. Four gene-modules related to ML with foamy microglia. **e,** Module eigengenes were correlated to lesion types. * q<0.1,** q<0.01, **** q<0.0001. **f,** Heatmaps depicting z-scored CPM values of genes in the selected modules. **g,** Selected hub genes for each module. **h-i**, Enriched cell types from brain or blood (**h**) and pathways (**i**) in the selected modules.

To identify the biological networks driving the foamy phenotype in an unbiased manner, we conducted a weighted gene co-expression network analysis (WGCNA) (Figure 3e-i, supp. files 1-3).^34^ We identified four modules—ME5, ME7, ME12, and ME15—that were significantly associated with MLs containing foamy microglia (Figure 3e). The microglia enriched ME12 module was most strongly linked to the foamy phenotype, while it was significantly downregulated in RL. This module featured *GPNMB*, *GRN* and *CHIT1* as hub genes and enriched for lysosomal genes (*PLA2G15, ACP5, ASAH1, DNASE2B, LGMN, CTSZ, CTSD*), antiviral response genes (*OAS1-3, OASL, ISG15, HLA-A, HLA-B, HLA-DQ, TRIM25, MX1, MX2*), and antigen presentation genes (*HLA-A, HLA-B, HLA-DQ*) (Figure 3i). The ME5 module, associated with microglia and other myeloid cells (Figure 3i), with hub genes *BTK* and *SYK*, consisted mainly of genes associated with damage-associated microglia (*SPP1, CLEC7A, CTSB, LPL,TREM2*), phagocytosis (*FCGR1A, FCGR3A, FCER1G, C1QA, MSR1, TREM2*), lipid metabolism (*PLIN2, PPARG, ALOX5, ALOX5AP, TBXAS1, APOC1, ACSL1*) and lysosomal processing (*CTSB*, *CTSS, CTSL, LYZ, VNN1, VAMP8, DPP4, TASL*). Module ME7 was related to the unfolded protein response and ER stress (*HSPA6, HSPH1*, *DNAJA1*, *XBP1*), while module ME15, containing *CD79A*, *CCL5*, and *IGHG1* as hub genes, linked to adaptive immune cells, particularly B- and plasma cells (Figure 3h). Given the general poor correlation between mRNA and protein levels, we investigated whether the gene-modules were confirmed at the protein level. Microglia-modules ME5 and ME12 were strongly replicated at the protein level, being significantly higher expressed in lesions with foamy microglia (Figure S10). Heat-shock protein module ME7 did, however, not replicate at the protein level. B-cell module ME15 was reflected in the immunoglobulin levels measured by proteomics (Figure S11). We found that predominantly immunoglobulin G1 (IgG1) increased in lesions compared to CWM, and IgG1 levels were significantly higher in ML with foamy compared to non-foamy (Figure S11c). In addition, we observed significantly higher expression of immunoglobulin receptors and complement components and their receptors in lesions with foamy microglia (Figure S12), indicating increased immunoglobulin mediated phagocytosis. In summary, WGCNA, validated by proteomics, reveals that ML with foamy microglia are characterized by an enhanced adaptive immune response, increased phagocytosis, lysosomal activity, lipid metabolism, and antigen presentation.

To assess the cell type composition of the lesions, we performed cell-type deconvolution of the bulk transcriptome using dTangle and a recently reported MS snRNAseq dataset.^22,35^ As expected, the number of oligodendrocytes was significantly lower in all lesions compared to CWM, while the number of astrocytes was increased (Figure S13). We also observed elevated B- and T-cell numbers, with B-cells being most abundant in foamy MLs (Figure S13c). This observation was confirmed by immunohistochemical analysis using the pan B-cell marker CD79A, showing a high abundance of B-cells in the perivascular space and infiltrating the parenchyma (Figure S11d-f). Microglia numbers increased across all lesions, with a tendency to be highest in MLs with foamy microglia (Figure S13c). However, this increase did not explain the gene expression differences between foamy and non-foamy samples, as most genes remained significantly different after correcting for microglia numbers (Figure S13e-f). This suggests that the observed WGCNA modules and differential expression are driven by gene expression specific to foamy microglia, rather than by a higher number of microglia.

Given the specific molecular profile of foamy lesions, we investigated whether the foamy phenotype represents a distinct microglial transcriptional state^36^. To this end, we examined snRNAseq-derived microglial states reported by Macnair *et al*. (Figure 4a), determined gene-signatures for each state and analysed their expression in the bulk transcriptome, adjusted for total microglia abundance (Figure 4b, S14). We found that the Micro_D profile was significantly elevated in lesions with foamy microglia compared to non-foamy, especially evident in MLs (Figure 4c,d). Conversely, the homeostatic cluster Micro_A was significantly reduced in lesions with foamy microglia but not in those with non-foamy microglia (Figure 4e). There was considerable overlap between the Micro_D profile and previously reported microglial states, such as the MIMS-foamy state reported by Absinta *et al.*^19^, amyloid-associated microglia in Alzheimer’s disease^37–39^, and lipid-associated macrophages (LAM) in obesity^40^ (Figure S15). The Micro_D profile was characterized by marker genes such as *GPNMB, CHIT1, APOC1, PLIN2*, and *ACP5* (Figure 4f,g, S16). Consistently, ontologies associated with the Micro_D state were lipid storage, phagocytosis, foam-cell differentiation, and lysosomal compartment (Figure 4h), but not pro-inflammatory signalling. This lack of classical pro-inflammatory signalling has been previously found in MS lesions by mass cytometry of microglia^41^ and matches the lipid-induced lysosomal phenotype described in other diseases^15,42^. Micro_D marker genes were also significantly enriched in lesions with foamy microglia at the protein level (Figure S17). *GPNMB,* a marker for lysosomal stress^43^, emerged as the strongest Micro_D marker gene (Figure 4f). Therefore, we performed double immunofluorescent staining on GPNMB to locate Micro_D microglia in MS lesions. GPNMB+ microglia were found in the rims of foamy MLs, but not in the lesion core. GPNMB positive cells were significantly higher in foamy MLs compared to non-foamy MLs (Figure 4i-l). This suggests that Micro_D GPNMB+ microglia, with increased phagocytosis, lipid metabolism and lysosomal stress, are involved in chronic lesion expansion.

**Fig. 4.**
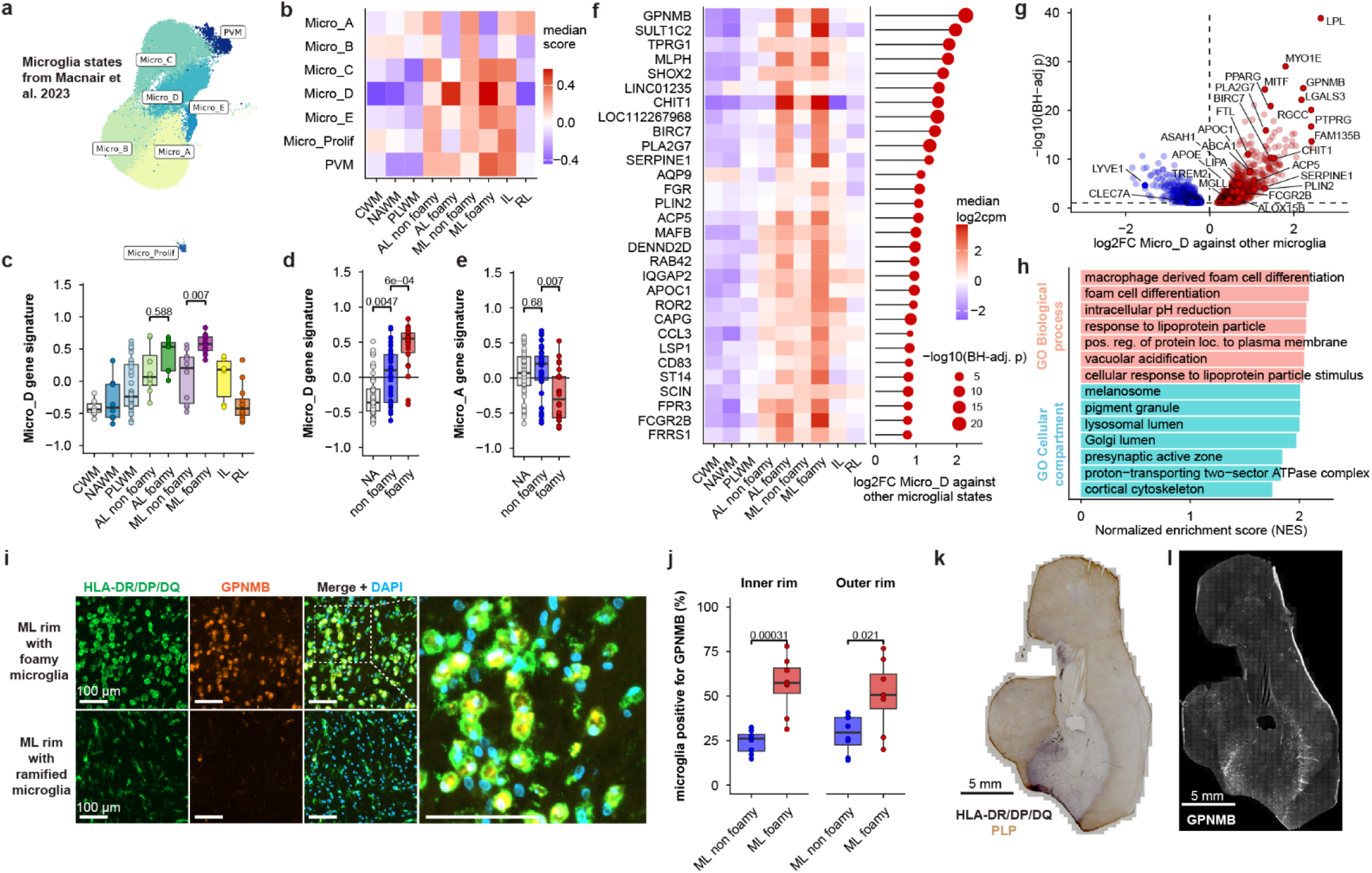
Foamy microglia represent a distinct microglial state. **a,** Microglial states by Macnair et al.^22^. **b-e,** Enrichment of the microglial state signatures from **a** in the bulk RNAseq data using gene set variation analysis (GSVA). P-values represent wilcoxon rank sum tests with BH-adjustment for multiple testing. **f,** Top 30 Micro_D marker genes. Markers genes were filtered for microglia-selective genes (>50% of counts originated from microglia). Left panel shows the median log2cpm in the bulk RNAseq dataset. The right panel shows the log2FC of micro_D marker genes against the combined other microglial states in the snRNAseq dataset. **g,** DEGs between Micro_D against the other microglial states calculated by limma and BH-corrected for multiple testing (FDR<10%). **h,** Pathways associated with Micro_D. **i,** GPNMB (orange) and HLA-DR/DP/DQ (green) double staining. **j,** Quantification of **i**. Inner rim indicates microglia within the demyelination border. Outer rim indicates outside the myelin border. p-values represent Wilcoxon rank sum tests. **k-l,** Adjacent sections from a representative ML with foamy microglia stained for HLA/PLP (**k**) or GPNMB (**l**).

Next, to obtain an integrated molecular profile that we could link to disease progression, we performed multi-omics factor analysis (MOFA) (Figure 5, S18)^44^. MOFA derives latent variables from covariance in the data, expressed as factors. We found that MS lesions in general, characterized by myelin loss, were captured by factors 1 and 2, whereas lesions containing foamy GPNMB+ microglia, represented by the Micro_D gene signature, were strongly associated with factor 3 (Figure 5b-e). Factor 3 contained increased cholesterol ester, BMP and oxylipin levels (Figure S19) along with their oxylipin-generating enzymes ALOX15B and TBXAS1, as well as IgG1, C1QC and *FCGR2B* (Figure 5h, S19), while axonal and synaptic proteins were downregulated (Figure S20)^16^.

**Fig. 5.**
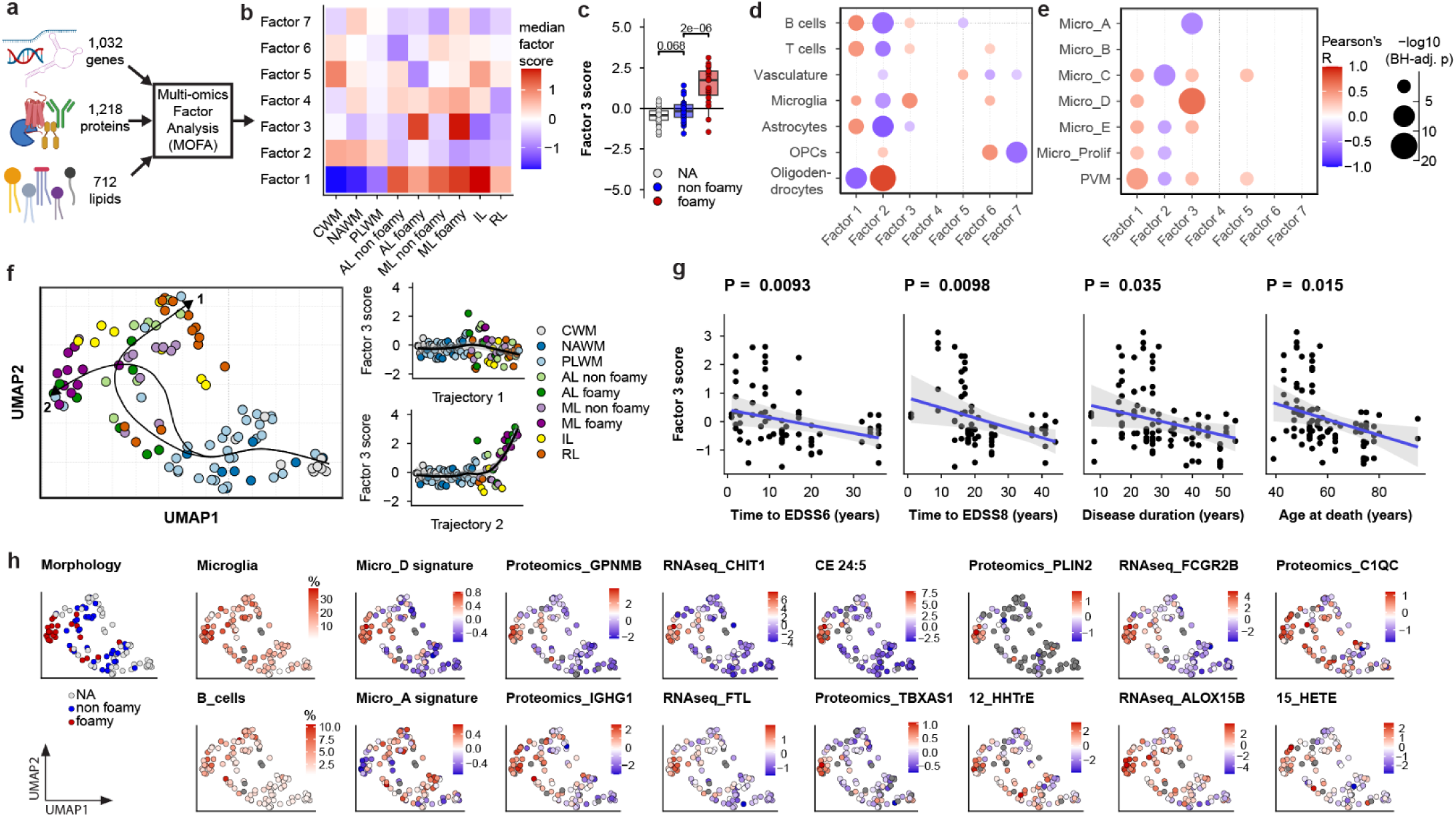
Multi-omics data integration reveals an integrated molecular profile of foamy microglia that associates with disease progression. **a-c,** Genes, proteins and lipids as input for multi-omics factor analysis (MOFA) delivered 7 factors based on a minimum 5% variance explained cutoff. Factor 3 significantly associates with foamy lesions (**c**, Wilcoxon rank sum test with BH correction). **d-e,** Correlation of MOFA factors with broad cell types (**d**) and specific microglial states (**e**). Data is presented as Pearson’s R, with size indicating BH-adjusted p-values. **f,** UMAP projection with a pseudotime trajectory analysis using the MOFA factors as input. Two trajectories were observed, differentiated by factor 3 scores. **g**, MOFA factor 3 was associated with clinical features in a generalized estimating equation (GEE) model to account for donor identity. **h,** UMAP projection as in **f,** but coloured by important variables that associated strongly with factor 3. For cell types, the colour depicts percentage, for Micro_A and Micro_D, it represents the GSVA score (from Fig. 4a**-e**). For all other plots each variable is expressed as the normalized log2-transformed value of its respective unit.

A slingshot pseudotime trajectory analysis^45^ revealed that factor 3 was a strong driver for the separation of remyelinated lesions from foamy MLs, while ALs and MLs with non-foamy microglia were more closely related to remyelinated lesions (Figure 5f). Notably, factor 3 also associated with more severe disease progression, as evidenced by shorter times to EDSS6 and EDSS8 and reduced disease duration (Figure 5g). Taken together, this indicates that foamy GPNMB+ microglia are a main source of variability in the molecular profile of MS lesions, which associates with more severe hallmarks of pathology and disease course. Mechanistically, our data suggests that sustained phagocytosis by the GPNMB+ microglia may overwhelm their endo-lysosomal pathway, leading to lipid droplet and oxylipin formation, which associates with enhanced adaptive immunity, complement and axonal damage that drives lesion expansion.

Finally, to identify potential drug targets to address the foamy MLs we performed activity-based protein profiling, a chemical biology technique to identify active enzymes in their native biological context (Figure S21)^46,47^. Because the foamy phenotype was characterised by increased lipid processing and oxylipin production, we performed ABPP using fluorophosphonate and β-lactone chemical probes targeting serine and cysteine hydrolases, which play a crucial role in lipid and protein metabolism^46,48^.

Among the 97 identified active hydrolases in MS lesions, several metabolic enzyme activities associated with MOFA factor 3 and were upregulated in foamy ML (Figure 6a-c). Notably, these included the lysosomal enzymes ASAH1, LIPA, CTSA, CTSG, CTSZ, PLA2G15 and PPT1, as well as PLA2G7, enzymes previously associated with the LAM^40^ and MIMS-foamy microglial state^19^. Notably, monoacylglycerol lipase (*MGLL)* activity, an enzyme associated with lipid droplets^49^, showed an approximately four-fold increase in foamy ML compared to non-foamy ML (Figure 6c), also significantly associated with MOFA factor 3 (Figure 6b), and its mRNA levels were highest in the Micro_D cluster (Figure S16f). MGLL catalyses the conversion of the endocannabinoid 2-arachidonoylglycerol (2-AG) into arachidonic acid (AA) in the brain, which is a precursor for oxylipins (Figure S22)^29,50^.

**Fig. 6.**
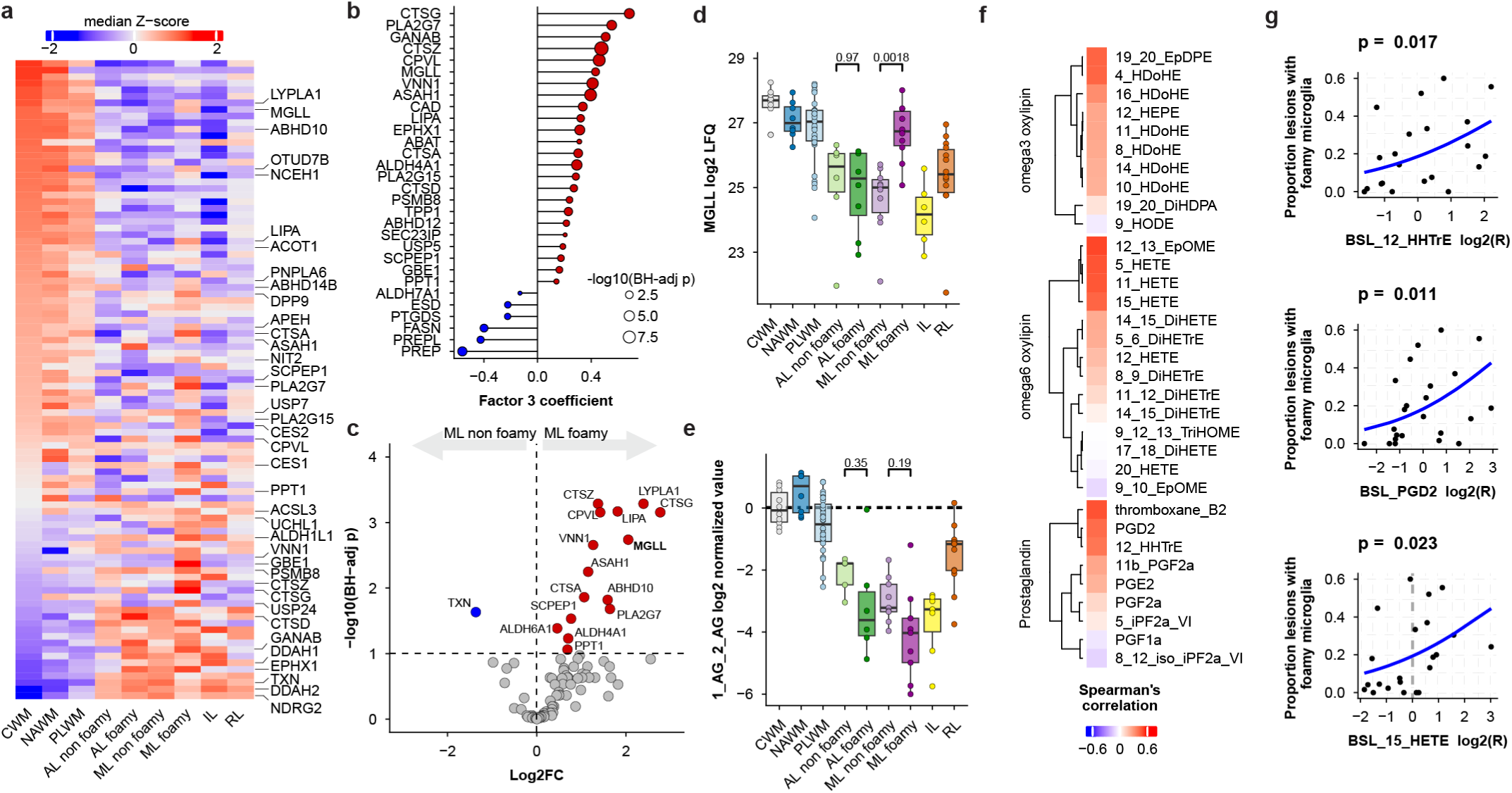
Foamy mixed lesions have high mono-acyl glycerol lipase (*MGLL*) activity and CSF oxylipins associate with higher foamy microglia load. **a,** Activity-based protein profiling detected 97 enzyme activities. The heatmap depicts median Z-scored LFQ values, sorted by CWM. **b,** Association of enzyme activities with MOFA factor 3. A linear model of the seven MOFA factors was fitted to the ABPP data, identifying enzymes associating to factor 3. **c,** Differentially active enzymes between MLs with foamy microglia to non-foamy. p-values were calculated by limma and BH-corrected for multiple testing (FDR<10%). **d,** MGLL activity across lesion types, p values are from **c. e**, MGLL-substrate 2-arachidonoyl glycerol (2-AG) levels across lesion types shows a trend for reduction in ML with foamy microglia compared to non-foamy. **f,** Spearman correlations of cerebrospinal fluid (CSF) oxylipin levels to the proportion of foamy lesions in patients. **g,** Linear models showing the significant association between selected oxylipins to the proportion of foamy lesions, p-values represent likelihood-ratio tests.

Because oxylipins and prostaglandins are excreted factors, we explored whether these lipids could be detected in the cerebrospinal fluid (CSF) of the same cohort of MS donors. CSF levels of many oxylipins positively correlated with the proportion of foamy lesions in their brain (Figure 6f,g). AA-derived oxylipins 12-HHTrE, PGD2 and 15-HETE, which were also strongly associated with MOFA factor 3 (Figure S23, 5h), exemplified this association, being significantly correlated to the proportion of foamy lesions. These results suggest that oxylipins produced by foamy microglia can be detected in the CSF.

Inhibition of MGLL increases 2-AG levels while decreasing free AA. Increased 2-AG levels affects neuroinflammation through activation of cannabinoid receptors CB1 and CB2^51,52^. This CB receptor activation reduces excitotoxicity^53,54^, dampens autoimmunity^55^, and shifts immune cells towards a pro-recovery, neuroprotective phenotype^56,57^. Additionally, the reduction in AA levels decreases the biosynthesis of oxylipins, which can drive neuroinflammation^50,58^. Indeed, MGLL inhibition has been shown to ameliorate EAE and cuprizone models of MS by reducing excitotoxicity, gliosis, and T-cell infiltration, while promoting remyelination^59–62^. Taken together, our data support a significant role for MGLL in human foamy white matter MS lesions, suggesting that its inhibition could not only provide symptomatic relief through cannabinoid receptors^63^, but also potentially serve as a disease-modifying therapy in secondary progressive MS by promoting lesion resolution and repair.

In conclusion, using a multi-omics approach, we have identified foamy GPNMB+ lipid-laden microglia as major drivers of variability in MS (mixed) active white matter lesions. This microglial phenotype is characterized by sustained phagocytosis and lipid accumulation, leading to lysosomal stress and oxylipin production. These lesions show high IgG1 levels, an increased number of plasma cells and elevated phagocytic receptor expression on microglial cells, suggesting that local antibody secretion may drive a hyper phagocytic phenotype, resulting in foamy microglia^64^. However, current B-cell depletion therapies do not resolve chronic lesion expansion^65^, indicating that CD20 directed depletion strategies do not address foamy MLs. This could be caused by ineffective clearing of parenchymal B-cells, or CD20-negative parenchymal antibody-secreting cells that are locally responsible for IgG1 production, further fuelling demyelination and lesion expansion^66,67^. BTK inhibitors, currently being tested in phase 3 clinical trials, could target foamy MLs as it is a core kinase controlling B-cell differentiation, as well as microglial activation^68^.

Additionally, oxylipins function as chemoattractants for adaptive immune cells, possibly further fuelling T- and B-cell infiltration^29^, enhancing Th17-cell^69^ and plasma cell differentiation and increasing immunoglobulin production^70–73^. These activities may create a feedback loop that perpetuates antibody secretion, microglial activation and further demyelination^29,63^. Disrupting this vicious cycle via MGLL inhibitors, currently tested in clinical trials, may offer an attractive therapeutic option for halting disease progression in secondary progressive MS. In addition, oxylipins, GPNMB or CHIT1^74^ in CSF or plasma^75^ may serve as potential biomarkers for patient stratification, assessing disease progression, and evaluating the efficacy of therapeutic interventions in MS progression. Finally, the foamy microglial state closely mirrors the microglia and macrophage phenotypes observed in several other diseases^15,36^, such as lysosomal storage disorders, atherosclerosis, Alzheimer’s disease and obesity, suggesting a possible common etiology in disease progression and therapeutic solutions.

## Supporting information

vlietetal_2024_supplementary figures

Supp_file_1_WGCNA modules genes with names and connectivity

Supp_file_2_Cell type enrichment for modules

Supp_file_3_WGCNA modules GO and KEGG terms

## Acknowledgements

We are grateful to the brain donors and their families for their commitment to the Netherlands Brain Bank donor program. Ahmed Mahfouz is kindly acknowledged for his advice in data analysis, particularly multi-omics integration. Bogdan I. Florea is acknowledged for managing proteomics facilities and equipment. Dennis Wever is acknowledged for his help in RNA isolation. TS and AJRH acknowledge support from NWO, funding the proteomics facilities through the X-omics Road Map program (project 184.034.019). DV and MvdS acknowledge funding from the Institute of Chemical Immunology. IH acknowledges funding from MS Research for funding the MS part of the Netherlands Brain Bank.

## Data availability

No custom software has been developed for the analyses described in this study. Raw RNAseq data will be available from gene expression omnibus (GEO) database. Mass spectrometry data will be available through the ProteomeXchange Consortium via PRIDE upon publication (Proteomics: PXD056856 and ABPP: PXD056899). All data processed data will also be available upon request from the authors. R code is available from the authors upon request.

## Author contributions

MvdS, IH, DvdV, LC and UG initiated the study. DvdV, AB, HJE, JQAC, IH & NBB contributed to donor selection, pathological characterization of the tissue, tissue dissemination and clinical data curation. DvdV isolated lesions and histologically verified lesion collection. -Omics data acquisition was performed by DvdV, BG (ABPP), XD, TH, ACH (lipidomics), TMS, AJRH (proteomics), DvdV (RNAseq) and AP (cytokine profiling). Data was analyzed by DvdV with contribution from TMS, XD, WM, MM, YZ, HJE and ED. DvdV, NvE, IACMV & CCB performed histological validation with immuno(fluorescent) stainings. Data was interpreted by DvdV, MvdS, IH, LC, XD, TMS, NvE, UG, ED, ME and AJRH. MvdS, IH, LC, UG and AJRH supervised the study. DvdV, MvdS and IH wrote the paper with input from all authors.

## Materials & Methods

### Human brain tissue

Human brain samples from 38 donors (28 Secondary Progressive MS and 10 matched non-demented controls) were obtained from the Netherlands Brain Bank (NBB, brainbank.nl, project No. 1327). All procedures of the NBB were approved by the ethical committee of Medical Ethical Committee of Amsterdam Academic Medical Centre and all donors gave written consent to the NBB for the use of their data and tissue for research. Fresh frozen tissue blocks containing sub-cortical white matter with and without MS lesions were dissected, snap-frozen in liquid nitrogen and stored at -80 °C. Mirrored pieces of this tissue were formalin fixed and paraffin embedded and stained to characterize different MS lesion stages (mixed active/inactive, active, inactive, remyelinated) [by the NBB](https://www.brainbank.nl/nbb-ms/characterization-of-ms-lesions/). Selection criteria for donors were a post-mortem delay (PMD, time-interval between the demise of the donor and freezing of the tissue) below 12 hours, a pH of the CSF higher than 5.5, and a clinician-confirmed diagnosis of secondary progressive MS for MS patients and absence of any neurological disease in controls, nor neurological disease-indicating pathology found in their brains as examined by a neuropathologist. The PMD ranged between 6:30 hours to 11:30, with a median of 8:45 hours. The pH of the CSF ranged from 5.8 to 6.8, with a median of 6.4. Donor characteristics were matched as closely as possible (Figure S1a), but there were differences in age between controls and MS donors.

### Analysis of lesion proportions and clinical features

Clinical data and pathological data was obtained from the NBB from MS brain donors that were collected by the NBB between 1990 and 2021 (N=250 MS donors). Time to EDSS6 and -8 were defined as the year in which EDSS6 or -8 was reached minus the year in which first symptoms were reported. Disease duration was defined as the number of years between first reported symptoms and death. Lesion counts were correlated to clinical features using a generalized linear model with a quasibinomial distribution of counts of lesions against other lesions, as previously described by Luchetti *et al*.^6^ P-values to test the model were calculated using the Likelihood-ratio test and corrected for multiple testing using the Bonferroni method.

### Lesion characterization

Lesions were classified using our previously defined criteria^6^ which is in line with the classification system by Kuhlmann and others.^1^ In short, control white matter (CWM) is white matter from control patients, absent of any sign of demyelination or aberrant microglial activation. Normal-appearing white matter (NAWM) is macroscopically intact white matter from an MS patient, where in the same tissue block there is no lesion or inflammation visible. Peri-lesional white matter (PLWM) is white matter adjacent to a white matter lesion. Depending on the lesion type the PLWM is adjacent to, sometimes there is considerable inflammation, or even the presence of foamy microglia. This was annotated as such. Active lesions (AL, type 2, also called acute) were defined as (partially) demyelinated areas with increased microglial presence throughout the lesion and no hypocellular core. Mixed active/inactive lesions (MLs, type 3, also called mixed, chronic active, smoldering or slowly expanding) were defined as a completely demyelinated areas with a hypocellular gliotic core, with an active rim of macrophages at the border of demyelination. Inactive lesions (IL, type 4) were characterized by complete demyelination, the absence of an active HLA^+^ rim and a hypocellular gliotic core. Remyelinated lesions (RL, type 6, also called shadow plaques) were characterized by partial myelination. In addition residual myelin should not only be myelin debris, but axonal myelin. Microglial activation was sometimes still present, but not to a larger density of cells than the surrounding PLWM. In addition to these classically defined lesion types, we profile the morphology of the microglia found in the active and mixed lesions.^6^ AL and ML with either foamy, or ramified and amoeboid (together called non-foamy) microglia were distinguished, in line with our previously published work^6,14^.

### Cryosectioning and isolation of MS lesions

Fresh frozen tissue was sectioned at 10 μm for stainings or 50 µm for tissue collection in a cryostat (Thermo Fisher HM525 NX) between -20 °C and -16 °C with Thermo Fisher MX35 Ultra microtome blades. Sections for stainings were mounted on Epredia Superfrost slides (J1800AMNZ), dried overnight using silica beads and subsequently sealed and stored at -80 °C until further use. Borders of a demyelinated lesion were identified based on Haematoxylin and Sudan Black staining and inspection under a light microscope. A surgical scalpel was used to demarcate the lesioned area, after which the lesion and peri-lesion were separated from 50 µm sections in a cryostat. Continuous stained sections were made to make sure the lesion borders were closely matched by the isolation method. Before and after every sample collected a slide was prepared for histological confirmation of lesion pathology. Tissue was collected in 1.5 mL sterile RNase-free SafeLock Eppendorf tubes, weighed and stored at -80 °C. Samples isolated for RNA extraction were stored in Trisure (BIO-38032, Bioline).

### Immunohistochemistry (single staining, CD79A, GFAP, CD3)

Frozen sections were taken from the -80 °C and warmed up to RT before opening the seal. The sections were then incubated in 4% PFA in PBS for 30 min, followed by peroxide blocking (1% H_2_O_2_, 0.5% Triton-X 100 in PBS) for 20 min. The section was subsequently incubated in primary antibodies (GFAP,1:1000, D1F4Q, Cell Signaling or CD79A, 1:200, M705001-2 DAKO or CD3, 1:100, A0452 DAKO), in BTHP-buffer (1% bovine serum albumin (BSA), 0.5% Triton-X100, 10% horse serum in PBS) overnight at 4 °C. Subsequently, secondary antibody was applied from the DAKO REAL EnVision Kit (K500711-2, DAKO) for 1 hour at RT. HRP immunoreactivity was then visualized with 3,3’-diaminobenzidine (DAB) using the same kit according to the manufacturer’s instructions (substrate was diluted 1:50 in buffer). After developing DAB staining, the sections were washed and stained in haematoxylin-solution (50 gr, L KAl(SO_4_), 1 gr, L haematoxylin, 0.2 gr, L NaIO_3_, 50 gr, L Chloral hydrate, 1 gr, L Citric acid). After washing, the sections were dehydrated in a dehydration series of 50% EtOH (3 min), 70% EtOH (3 min), 96% EtOH (5 min), 100% EtOH (5 min), 100% EtOH (5 min), xylene (10 min) and xylene again (10 min). After the last xylene step, the section were dried and mounted in Entellan (1079600500, Merck Millipore) using 24×50mm coverslips (CLS2975245, Corning). Sections were imaged on a Zeiss Axioscan 7 and further processed using Qupath software (v0.5.0).^76^

### Immunohistochemistry (double staining, HLA+PLP)

Sections were fixed and peroxide blocked as described above. Sections were subsequently incubated in primary antibody against HLA-DR, DQ, DP (1:1000, M0775, DAKO) in BTHP-buffer (1% BSA, 0.5% Triton-X100, 10% horse serum in PBS) for 1 hour at RT. Subsequently, secondary antibody was applied from the DAKO REAL EnVision Kit (K500711-2, DAKO) for 1 hour at RT. HRP immunoreactivity was then visualized using DAB, Ni^2+^ staining (0.5 mg/mL DAB tetrahydrochloride, 2 mg/mL nickel ammonium sulfate, 0.01% H_2_O_2_ in PBS) for ∼10 min. Subsequently, a primary antibody against PLP was incubated on the section (1:1000, clone plpc1, MCA839G, Bio-Rad) in BTHT-buffer (1% BSA, 0.5% Triton-X100, 10% Horse serum in TBS) overnight at 4 °C. The next day, the secondary antibody from the DAKO REAL EnVision Kit was applied again for 1 hour at RT, before developing normal DAB staining using the same kit according to the manufacturer’s instructions (substrate was diluted 1:50 in buffer). After developing DAB staining for PLP, the sections were washed, stained in Haematoxylin-solution, mounted and imaged as described for the single staining.

### Immunofluorescent analysis of GPNMB positive cells

Frozen sections were taken from the -80 °C and warmed up to RT before opening the seal, or freshly sectioned on the cryostat. The sections were then incubated in 4% PFA in PBS for 30 min, followed by blocking (10% Donkey serum, 0.5% Triton-X 100 in PBS) for 2 hours at RT. The section was subsequently incubated with primary antibodies (GPNMB,1:500, E4D7P, Cell Signaling & HLA-DR/DQ/DP, 1:1000, M0775, DAKO), in 0.5% Triton-X100, 5% Donkey serum in PBS, overnight at 4 °C. Appropriate secondary antibodies (Jackson Immuno, Donkey anti-IgG (H+L), 1:1000) were incubated subsequently for 2 hours at RT in PBS with 0.5% triton-X. Autofluorescence was quenched using by incubating the section in 0.5% Sudan Black B in 70% Ethanol for 2 min. subsequently, the sections were briefly rinsed in 50% ethanol and subsequently with PBS. Nuclei were stained with 1 µg/mL DAPI (Sigma) and the sections were mounted under a coverglass (Corning) in Mowiol 4-88 mounting medium (100 mg/mL Mowiol 4-88, 25% glycerol, 25 mg/mL DABCO in 0.1M Tris-HCl pH8.5).

Images were captured using a Axioscan 7 slidescanner (Zeiss) and quantification of positive cells in each lesion was performed using using QuPath (v0.5.0). Regions of interest were selected using the wand- and brush annotation tool, based on demyelination and the border of microglia/macrophages in the HLA/PLP staining. The region where microglia/macrophages were located in the demyelinated part of the rim was called the ‘inner rim’. The region where the microglia/macrophages were located in the myelinated part of the rim was called the ‘outer rim’. After region selection in each lesion, analysis of HLA and GPNMB was performed using *Cell detection* and *Object classifier* commands. Cell detections were performed within the region of interest using DAPI staining to detect the nucleus of individual cells combined with the cell expansion option to segment as much surface area of the whole cell as possible. A Random trees (RTrees) classifier was trained to separate microglia/macrophages from other detections. Objects were filtered for ‘Cells’ and all ‘’AF488’’ features were selected. Training images were created by creating duplicate channel training images for AF488. The object classifier was trained on ‘Points only’ on two classes, “AF488” and “ignore*”. Approximately 50 points for each class were annotated by hand in 6 training images of each lesion type (gives a total of 100 points per image in 12 training images). A second, separate, Random trees (RTrees) classifier was trained to separate GPNMB-positive cells from other detections. This was done in the same way as the classifier for microglia/macrophages, with all ‘’AF555’’ features selected. The classifiers were loaded and applied sequentially by selecting them both in the option *Load object classifier*. The measurements were exported and further analysed in R.

### Cytokine profiling

Cytokine concentrations were measured using a custom-made Human Luminex Discovery Assay (LXSAHM-26) according to the manufacturer’s protocol. Lysates were prepared from whole lesions, white matter by glass-bead mechanical lysis in 20 mM HEPES, 1 mM MgCl_2_, 2 U/mL benzonase. After protein quantification using the Bradford method (Bio-Rad), lysates were snap-frozen in LN_2_ and stored at -80 ⁰C until further use. 25 µL of 1 mg/mL lysate was thawed and diluted 1:2 in calibrator diluent buffer (RD6-52). The standard cocktails were reconstituted in RD6-52 buffer and further diluted 1:3 following the manufacturer’s instructions. In total, 50 µL of sample or standard were mixed with 50 µL of diluted microparticles cocktail and incubated for 2 hours at room temperature on a horizontal orbital microplate shaker operating at 800 rpm. After the incubation, the plate was washed three times with a magnetic plate washer, using 150 µL of wash buffer and allowing one minute before removing the liquid. Next, 50 µL of diluted Biotin-Antibody Cocktail were added to each well and the plate was incubated for 1 hour at room temperature while shaking at 800 rpm, followed by another washing step and the addition of 50 µL of diluted Streptavidin-PE. After 30 minutes incubation and a washing step, the microparticles were resuspended in 100 µL wash buffer. The plate was analysed with FLEXMAP 3D using Standard PTM settings and doublet discriminator gates set at 8000 and 16500. After assigning the microparticle region for each measured analytes, 50 µL samples were acquired. Cytokine concentrations were quantified by the instrument software (BioPlex Manager) using a five-parameter logistic (5-PL) curve.

### Proteomics sample preparation

Lysates were prepared from tissues by glass-bead mechanical lysis in 20 mM HEPES, 1 mM MgCl_2_, 2 U/mL benzonase. After protein quantification using the Bradford assay (Bio-Rad), lysates were snap-frozen in LN_2_ and stored at -80 ⁰C until further use. Three µg of protein was resuspended in 20 µl of SDC buffer (2% sodium deoxycholate (SDC), 10 mM tris(2-carboxyethyl) phosphine (TCEP), 10 mM Tris (pH 8.5), 40 mM chloroacetamide) supplemented with Complete mini EDTA-free protease inhibitor cocktail (Roche). The samples were denatured at 95°C for 5 min. Cooled, reduced and alkylated samples were mixed with 162 µl of 50 mM ammonium bicarbonate, followed by the addition of trypsin (Promega) and LysC (Wako) in 1:50 and 1:75 ratios respectively. The digestion took place overnight at 37°C. The digestion was quenched with formic acid at the final concentration of 2% and centrifuged at 20 000 g for 20 min to remove the SDC. The acidified supernatant containing 800 ng of peptides was loaded onto the EvoSep StageTips (EV2018) according to the manufacturer’s protocol.

### LC-MS/MS analysis proteomics

The samples were analyzed on EvoSep One liquid chromatography system (EvoSep, Denmark) coupled to Exploris 480 mass spectrometer (Thermo Fisher Scientific, Bremen). The peptides were eluted from the EvoSep StageTips and separated on an EvoSep analytical column (15cmx150 µm, 1.9 µm; EV-1106) using a 44 minute gradient (30 SPD program) and the data was acquired in data-independent mode. The following mass spectrometric parameters were used for the full MS scan: scan mass range set to 375-1100 *m, z*, resolution of 60 000 at 200 *m, z*, AGC target set to standard, maximum injection time set to auto. For the MS, MS spectra the following parameters were used: quadrupole isolation window of 15 Da with 1 Da overlap between the windows (total of 40 windows), the precursor mass range was set to 400-1000 *m, z*, resolution of 15 000 at 200 *m, z*, normalized AGC target was set to 1000%, injection time set to auto, normalized collisional energy was set to 27%, isotope exclusion set to On.

### Raw data processing proteomics

All raw files were processed with DIA-NN software (version 1.8.1) with the deep learning *in silico* spectral library generation option. The digestion was set to trypsin with 1 missed cleavage allowed. Cysteine carbamidomethylation was set as a fixed modification and methionine oxidation was set as a variable modification. The threshold FDR for the protein identification was set to 1%. The full scan mass accuracy was 6 ppm and the optimized MS, MS mass accuracy was set to 22 ppm. All other settings were set to default. The uniprot human database with 20398 entries was used (release April 2023) for the search. The quantification was based on the unique peptides for the downstream analysis. Two samples were excluded from the downstream analysis as they contained a too high percentage of missing values.

All the downstream analyses were performed in R (v4.4.1). Proteins that had at an LFQ value in at least 70% (found, total ≥ 0.7) of samples in one group (lesion type) were included, and immunoglobulin variable regions were excluded. This led to 3,237 proteins included in downstream analysis. Missing values were imputed by sampling from a normal distribution around the lowest value found for a given protein, with a standard deviation (sd) of 1/3 of the original sd. The imputed dataset was only used for principal component analysis, all other analyses were performed with the original dataset without data imputation.

### RNA extraction

Isolated lesions were dissolved in 500 µL Trisure (BIO-38032, Bioline). Subsequently, 100 µL CHCl_3_ was added to the tube and the solution was centrifuged (15 min, 11,000 rcf, 4 ⁰C). The aqueous layer was carefully removed and 500µL was added to the same tube for an additional extraction. After combining the two aqueous layers 1 equal volume 70% EtOH was added and the solution was added to an RNAeasy column (QIAgen). The RNA was washed with RW1 buffer, after which any DNA contamination was removed by incubation with DNase (15 min, RT). After this the column was washed in subsequent steps with RW1 buffer, then with RPE buffer and lastly with 80% EtOH. The column was air dried for 5 minutes before eluting the RNA in 16 µL RNAse free water. 1 µL of the extracted RNA was used for analysis on a Denovix DS-11 spectrofotometer to analyze the purity and concentration.

### RNA sequencing and analysis

Isolated RNA was sent for total RNA sequencing at Genomescan (Leiden, The Netherlands). RNA integrity was assessed, delivering an RNA quality value (RQN) value, which is used in downstream analysis as a covariate. No minimum RQN value was set as a requirement. After library prep, a library QC was performed which gave satisfactory results. Subsequently, the libraries were sequenced on a NovaSeq 6000 sequencer for a target library size of 30 million reads. Alignment was performed using HISAT v2.2.0 using reference Ensembl genome human GrCh 38. Ensembl identifiers were converted into gene names using AnnotationDbi^77^ and the human reference genome database Org.Hs.eg.db (v3.16.0)^78^ from Bioconductor.

The raw count data were subsequently loaded in R (v4.4.1) and a DGEList object was created using EdgeR. Samples with library sizes < 2 million were excluded. This threshold was determined by inspecting count distributions across the samples. These criteria led to the exclusion of four samples, leaving 105 samples for subsequent analysis. Count distributions were found to be satisfactory and mostly uniform after the exclusion of the 4 samples with low library sizes.

Criteria for gene filtering (n=61,541) were having > 2 counts per million (CPM) on average in at least one sample type using the ‘cpmbygroup’ function in EdgeR. Additionally, we excluded immunoglobulin variable regions and only kept constant regions. This left 16,652 genes for further analysis. CPM values were calculated using log2(counts+0.5 per million) with normalization using the trimmed-mean method (TMM) in EdgeR. These data were inspected for correlation with technical and biological covariates using principal component analysis (PCA). CPM values used in PCA analysis, MOFA and WGCNA and all heatmap visualizations were corrected for covariates using the ‘RemoveBatchEffect’ function in *Limma*, with the design *y = ∼0 + Lesion type + RQN + sex*, Lesion type categories (9 groups) being the design, and RQN and sex being the covariates to be regressed out.

### Weighted gene coexpression network analysis (WGCNA)

A WGCNA-based co-expression network was constructed using the WGCNA package (v1.72-5), and its dependencies fastcluster (v1.2.6), dynamicTreeCut (v1.63-1), preprocessCore (v1.66.0 and impute (v1.78.0). The transcriptomics input was corrected for *RQN* and sex as described above. Because we were interested in the differences between lesions and not necessarily between lesions and control or normal appearing white matter, we excluded all non-lesion samples. Indeed, a constructed network using all samples dissected the data into fewer modules, with two main modules that represent genes upregulated in all lesions, and genes downregulated in all lesions (data not shown). Genes for WGCNA input were selected for high variance, including those genes with a standard deviation within lesion samples higher than the mean standard deviation, yielding an input dataset of 52 lesion samples and 6,547 genes.

To determine the power threshold β, the function PickSoftThreshold was used, where we tested powers between 1 and 20, and then assessed scale-free topology. At β = 13, the threshold (R^2^ between log(freq) and log(connectivity) = 0.85) for approximate scale-free topology was reached. Next, WGCNA Modules were generated using the BlockwiseModules function, with corType set to “bicor”, TOMtype set to “signed”, networkType set to “signed”, a power β of 13, mergeCutHeight set to 0.2 and a minimum module size of 30. A signed network was chosen to preserve the directionality of the generated modules, improving interpretability of GO terms associated with the module. The biweight midcorrelation (“bicor”) was chosen over the Pearson correlation to make the method less sensitive to outliers and more robust, which generates more meaningful modules, as the authors of WGCNA recommend^34^.

This dissected the dataset into 16 modules ranging in size from 56 to 968 genes. 435 genes were not assigned to any module (Module 0). Module eigengenes were calculated using the function ‘moduleEigengenes’. The module eigengenes consistently explained high percentage of variation within the module (except for Module 0), ranging from 38.5% to 59.2% indicating that these eigengenes represent the modules well.

Module eigengenes were correlated to traits, in this case the lesion types, using the pearson correlation, and with subsequent calculation of p-values by the function ‘corPvalueStudent’. Acquired p-values were corrected for multiple testing of the 17 modules using the Benjamini-Hochberg method with an FDR set at 10%.

GO terms associated with module-genes were found using clusterProfiler (v4.6.1) and the org.Hs.eg.db_3.19.1. We used the enrichGO function with all WGCNA input genes as background (universe), the module genes as input, ontology set to “ALL”, minGSSize set to 10 and MaxGSSize set to 500, and a p-value cutoff of 0.05. KEGG pathways were found using the enrichKEGG function, with the same inputs. Heatmaps showing expression data for the modules were generated using the ComplexHeatmap package^79^ (v2.20.0) with the module genes log2 Z-scored counts per million as input.

Cell-type enrichment was performed using the function *UserListEnrichment* with a reference list of biomarkers of various cell types from CellMarker 2.0, a database containing an up-to-date manually curated collection of markers of various cell types in different human tissues^80^. All the markers from brain and blood were extracted from the dataset and pre-processed into an input reference list including 915 categories (Cell types) and 6517 markers (genes). All modules were tested for cell type enrichment, except module 0. P-values were corrected with Bonferroni multiple testing correction, which is default in the WGCNA workflow and *UserListEnrichment* function.

### Statistics and differential expression

No power analysis was performed to determine sample sizes. All differential expression/abundance analyses were performed using the Limma linear modeling framework^81^. For the RNAseq data we constructed a linear model taking into account covariates, using the formula *y = ∼0 + Lesion type + RQN + sex, w*ith Lesion type being the design and RQN and sex being covariates. We ran Limma’s linear modelling function ‘lmFit’ using the modeling design as above, using *voom* for precision weight analysis.^82^ For all other datasets we ran the lmFit function on log2-expression values without *voom* and without taking any covariates along in the model (y= ∼0 + Lesion type).

For calculating differential expression between foamy and non-foamy microglia, which can appear in several backgrounds (mixed lesions, active lesions and also peri-lesional white matter), we constructed a linear model using lesion type as a covariate, without specifying the morphology (7 groups: CWM, NAWM, PLWM, AL, ML, IL, RL) and specifying the morphology as separate parameters. For the RNAseq the linear model was y = ∼0 + morphology + Lesion type (7 groups) + RQN + sex and for all the other datasets y = ∼0 + morphology + Lesion type (7 groups).

These linear models were fed into the limma eBayes algorithm with default settings. Log2-fold changes and Benjamini-Hochberg corrected p-values were extracted using the TopTable function. We generally accept a false discovery rate (FDR) of 10%, meaning that we consider variables with a BH-adjusted p-value lower than 0.1 statistically significant.

Overlap between sets of differentially expressed genes compared to NAWM was visualized using the upset-plot function from ComplexHeatmap (v.2.20.0).^79^

### Principal component analysis (PCA)

PCA was performed using the ‘prcomp’ function in base R. ‘scale’ was set to FALSE, to preserve highly variant variables to have more weight in the PCA. PCA was performed on the full proteomics, lipidomics and ABPP datasets. For RNAseq PCA was performed on the 1,000 most variable genes. The obtained principal components were rotated to focus loadings of variables onto a single component using the ‘varimax’ function. Correlation of principal components to metadata was performed using the PCAtools package, using the function ‘eigencorplot’. The number of components was determined using the elbow criterion, in PCAtools automated in the function ‘getelbow’.

### Multi-omics factor analysis (MOFA)

MOFA was performed based on the workflow and code by Argelaguet et al.^44^ using the R package MOFA2 (v1.8.0). A python dependency was established using Basilisk. Data input for MOFA were the full lipidomics dataset (712 lipids) and a reduced RNAseq and proteomics dataset. Only highly variable transcripts were selected by filtering for genes with an sd higher than 2*mean(sd) of the full the dataset, which yielded 1,032 transcripts. Proteins were selected by filtering for proteins with an sd higher than 1*mean(sd) of the full the dataset, which yielded 1,218 proteins. 92 samples had data for all three data modalities. 11 samples had data for two data modalities and 7 samples had data from only one data modality (6 RNAseq, 1 proteomics).

We ran the MOFA model with largely default settings. Scale_views was set to ‘TRUE’, convergence mode was ‘medium’ and maximum iterations was 2000. We obtained 7 factors that explained a minimum of 5% variance in at least one data modality. The total variance explained by 7 factors were 53.5% for the RNAseq input, 57.7% for the proteomics input and 72.8% for the lipidomics dataset. Factor values were tested for differential expression across groups using wilcoxon rank sum test using the rstatix package (v.0.7.2), with correction for multiple testing using Benjamini-Hochberg FDR.

Dimensionality was further reduced using uniform manifold approximation and projection (UMAP) using the package uwot (v.0.1.16) using a n_neighbours parameter of 20. Pseudotime trajectories were calculated using Slingshot (v2.6.0) using clusters generated by Mclust (v6.1) using a G parameter of 5. The starting cluster was defined as the cluster containing control white matter samples. Generated trajectories were extracted using the ‘slingCurves’ function and plotted onto the UMAP projection using ‘geom_path’. Trajectories were plotted against the original input data (factor values) and a smooth fit was plotted using the ‘loess’ fitting function of ggplot2.

MOFA factor 3 was associated with the clinical features in a generalized estimating equation (GEE) model to account for donor identity, because our samples were nested within donors (100 samples from 28 MS donors). To this end, we used the function ‘geeglm’ from the package geepack (v1.3.10), with a gaussian distribution and an exchangeable correlation structure.

### Gene set enrichment analysis (GSEA)

GSEA was performed using log2-fold changes as input with the ClusterProfiler package (v4.6.2), function ‘gseGO’. Ontology was set to “ALL”, the database was Org.Hs.eg.db (v3.16.0), maxGSSize=100, minGSsize=10, eps=1e-300 and pvaluecutoff=0.05. MOFA factor loadings from proteomics and RNAseq were also investigated using GSEA, using the same settings, with the relative factor loadings (from -1 to 1) as input. Lipid class enrichment was performed using fgsea (v1.24.0) using custom lipid class lists (31 classes).

### snRNAseq dataset processing & deconvolution

Single-nucleus RNA sequencing data was obtained from Macnair et al.^22^ White matter samples were selected and counts from the same broad cell type (7 cell types) were combined into one pseudobulk per sample. Then, a DGEList object was created using EdgeR and the samples were normalized using the ‘TMMwsp’ method in EdgeR. This was chosen over normal TMM, because the snRNAseq data contains many zero’s which affects normalization. Samples were then filtered for minimum library size of 20,000. Genes were filtered for highly expressed genes by requiring a mean CPM in at least one cell type of 10 using the ‘cpmbygroup’ function. This gave a dataset of 533 pseudobulk profiles across 7 cell types (astrocytes, oligodendrocytes, OPCs, microglia, vasculature, B_cells and T_cells) with 15,084 genes.

Marker genes were calculated using limma, where we tested each celltype against the combined other cell types. Differentially expressed genes were determined using voom and eBayes functions as described above. Further criteria for marker genes were a log2FC of at least 4 against the average of the other cell types, an adjusted p-value of smaller than 0.00001, and that the marker gene was unique. We then restricted the list to the top 100 best marker genes (ranked by logFC*-log10(adjusted p-value)) for deconvolution. For B cells, we excluded immunoglobulin genes from the analysis, because their expression per B-cell is extremely variable and can be extremely high, leading to overestimation of the number of B-cells. Given these criteria, not every cell type had 100 marker genes. The number of marker genes were between 75 and 100. Plotting these marker genes against the pseudobulk profiles consistently gave very clean profiles (Figure S7a). Deconvolution was subsequently performed using the R package ‘dtangle’ (version 2.0.9). The input data for dtangle were:

1. The log2(cpm) bulk data normalized by edgeR without any corrections for covariates.
2. The log2(cpm) pseudobulk data from snRNAseq normalized by edgeR as described above.
3. The lists of marker genes

Further settings were a gamma of 0.9433902 as recommended by the authors of dtangle for RNAseq data, and the option “summary_fn” was set to “mean”.

To validate the method and the marker genes, the pseudobulk dataset was randomly split into two parts (80% training, 20% test). We then used the dTangle algorithm with the same settings, and the same marker genes to predict the cell type composition of the pure pseudobulk profiles in the test dataset. This consistently gave very high percentages of the correct cell type around ∼90% (Figure S13b).

### Evaluation of microglia subclusters

Microglial states were analyzed from the single-nucleus RNA sequencing data from Macnair et al 2023.^22^ White matter samples were selected and counts from microglia subclusters (7 microglia clusters: Micro_A, Micro_B, Micro_C, Micro_D, Micro_E, PVM, Micro_Prolif) were combined into pseudobulk per sample. Then, a DGEList object was created using EdgeR and the samples were normalized using the ‘TMMwsp’ method in EdgeR. Pseudobulk profiles were then filtered for minimum library size of 10,000. Genes were filtered for highly expressed genes by requiring a mean CPM in at least one subcluster of 10 using the ‘cpmbygroup’ function. This gave a dataset of 442 pseudobulk profiles across 7 subclusters with 11,575 genes.

Marker genes for each microglia subcluster were calculated using using limma, where we tested each microglia subcluster against the combined others. Differentially expressed genes were determined using voom and eBayes functions as described above. Further criteria for marker genes were a positive FC, a BH-adjusted p-value of smaller than 0.05, and that the gene was expressed mainly in microglia (more than 50% of all counts in the complete snRNAseq dataset came from microglia). Given these criteria, we identified between 24 and 328 marker genes for each subcluster. The marker genes were ranked based on logFC, and the top 30 markers genes were selected to compose a gene-signature. To then estimate the relative presence of each microglial subcluster in the bulk RNAseq dataset, we used gene-set variation analysis (GSVA)^83^ with each marker gene list as a gene-set. To control for microglial numbers in the bulk RNAseq data, we took the estimated microglial proportion as described above, and we used Limma’s ‘RemoveBatchEffect’ function to correct for the log2-transformed microglia abundance, as well as sex and RQN. This corrected dataset was used in GSVA. Further settings for GSVA were kcdf set to ‘Gaussian’ and maxDiff set to ‘TRUE’. This resulted in a GSVA score per microglial state between -1 and 1 for all samples. We then tested all microglia subcluster signatures between lesion types, and between foamy and non-foamy samples using the Wilcoxon-rank sum test with Benjamini-Hochberg correction for multiple testing.

Deconvolution of the microglia states was also tried using dtangle as described for the broad cell types. However, this did not yield good results because there is too much collinearity between the distinct microglia subclusters. Validation of this method using a part of the snRNAseq pseudobulk did not purely enrich the right cell type, but also enriched other microglial subclusters (data not shown).

### Lipidomics sample preparation

Samples (50 µm sections) were lysed, and protein concentrations (Bradford method) were quantified. Two study QC pools (Normal white matter pool and lesion pool) were prepared using 10 - 100 μL of more than 50% of all the samples. 11 QC aliquots were made separately from the normal white matter pool and lesion pool.

Purchased or synthesized standards, internal standards (IS) were dissolved in methanol, ethanol, chloroform or ACN. These stock solutions were further diluted and mixed to make the standard stock solutions and IS stock solutions. The eCB synthesis IS mix contained 18:1, 18:1-PE-N-17:0, p18:1, 18:1-PE-N-17:0, 18:1-OH-PE-N-17:0, p18:1-OH-PE-N-17:0, OH-OH-PE-N-17:0. The lipids IS mix contained deuterated version of ceramides, (lyso)phospholipids, diacylglycerols, triacylglycerols and cholesterol esters. The signaling IS mix contained deuterated version of oxylipins, endocannabinoids (eCBs), free fatty acids and bile acids. The eCB synthesis calibration standards contained standards of precursors of eCBs, the signaling calibration standards contained oxylipins, eCBs, free fatty acids and bile acids, the oxidized lipids calibration standards contained oxidized versions of phosphatidylcholines. The IS mixes and calibration standards were stored at -80 °C. The mixed IS working solutions were prepared and stored at -20 °C till usage.

Aliquots of sample lysates (∼1mg protein/mL, 50 μL) were thawed on ice in 1.5 mL Eppendorf tubes. To each sample 10 μL IS work solution was added. Calibration samples were prepared by spiking 10 μL of each calibration standards into 50 μL of water. Extraction was performed by 100 μL extraction buffer (0.2 mM Ammonium formate) and 1000 μL extractant (MTBE:BuOH, 50:50, v/v). Samples were then mixed in a Next Advance Bullet Blender (5 min, 90% speed, room tempreture) followed by centrifugation (16,000 x g, 10 min, 4°C). 950 μL of the organic layer was transferred into clean, pre-cooled tubes and concentrated in a SpeedVac vacuum concentrator (ThermoFisher Scientific, Waltham, USA), followed by adding 50 µL of reconstitution solution (MeOH:ACN, 30:70, v/v) and agitating for 15 min. The reconstituted samples were centrifuged (16,000 g, 10 min, 4°C) and 40 µL were transferred into autosampler vials with inserts. Samples were kept at -80 °C till LC-MS analysis.

Samples were randomized and run in 1 batch. Each batch included QC samples and blank samples. QC samples are used to assess data quality. Each batch included study QC samples. Method blanks (proc blanks) are used to check for background signal. These samples are composed of analyte-free matrix and have gone through all the steps of the sample preparation procedure with the reagents only.

### CSF lipidomics sample preparation

300 µL of fresh frozen cerebrospinal fluid was spiked with 10 µL IS calibration standard, and subsequently extracted using the same method as described above. For the CSF lipids, only the signaling lipids platform was measured.

### Signaling lipids platform (oxylipins, endocannabinoids)

The signaling lipids platform covers 196 metabolites, including various isoprostane classes together with their respective prostaglandin isomers from different poly unsaturated fatty acids (PUFA), including n-6 and n-3 PUFAs such as dihomo-γ-linoleic acid (DGLA) and arachidonic acid (n-6) and eicosapentaenoic acid (EPA) and docosahexaenoic acid (DHA) (n-3). Endocannabinoids, endocannabinoid analogues and bile acids are also included in this platform. Reference standards were used for each analyte for peak identification, 47 deuterated internal standards were used for the correction of variations from sample preparation and LC-MS runs^16^.

A QTRAP 7500 (AB Sciex, Concord, ON, Canada) was coupled to an Exion LC AD (AB Sciex, Concord, ON, Canada). MS, MS experiments were done with a Turbo V source (AB Sciex, Concord, ON, Canada) operated with ESI probe. An Acquity UPLC BEH C18 column (Waters) was used to measure the samples. The three-pump LC system consisted of mobile phase A (MPA, H_2_O with 0.1 % acetic acid), mobile phase B (MPB, 90 % CH_3_CN, 10 % MeOH with 0.1 % acetic acid) and mobile phase C (MPC, IPA with 0.1 % acetic acid). The injection volume was 5 μL stacked with 10 μL of mobile phase A. The flow rate was 0.7 mL, min, and each run takes 16 min. The gradient started at 20 % MPB and 1% MPC. The MPB progressed from 20 % to 85 % between 0.75 min and 14 min while the MPC ascended from 1 % to 15 % between 11 min to 14 min after which conditions were kept for 0.3 min and then the column was re-equilibrated at initial conditions until 16 min. An electrospray ionization source (ESI) was used with parameters: interface temperature 600 °C, curtain gas 45 psi, CAD gas 9 psi, gas 1 and gas2 both 65 psi. The mass spectrometer operated in polarity switching mode and all analytes were monitored in dMRM mode. Data was acquired using Sciex OS Software V2.0.0.45330 (AB Sciex)^16^. Assigned MRM peaks from the acquired data were integrated using SCIEX OS (version 2.1.6) Software and signals were corrected using proper internal standards.

Blank effects (BE) for each analyte were checked by comparing proc blank samples to quality control (QC) samples. The precision and reproducibility of the analytical process were checked using the relative standard deviations (RSDs) of the QCs.

51 analytes complied with the high-confidence acceptance criteria of QC RSDs ≤ 15% and BE ≤ 40% in both QC pools (Normal white matter pool and lesion pool). 11 analytes had one or two of the QC RSDs within 15-30.6% or BE > 40% in one or both QC pools. These metabolites with high blank effect (>40%) were also reported as we know the blank effects are from the extractant, which is consistent and under control levels. The data is reported as relative response ratios (target area, ISTD area; unit free).

### HILIC MS/MS based lipids platform (phospholipids, sphingolipids, BMPs)

This forward phase Lipid platform covers 1320 nonpolar lipids targets. The samples were measured using 3 separate acquisition methods. Acquisition method 1 measurement contains classes of phosphatidylcholine, phosphatidylinositol, phosphatidylserine, phosphatidylglycerol, and bis(monoacylglycerol)phosphates. Acquisition method 2 measurement contains sphingomyelins, hexosylceramides, lactosylceramides and phosphatidylethanolamines. The acquisition method 3 measurement contains triglycerides. The identification of metabolites in this platform are based on the dimensions of specific MS, MS transitions and retention time (RT), Using HILIC columns, the lipids from the same class elute in a narrow RT window, while different lipid classes elute at different RTs. For each class, one or more deuterated internal standards were used to check the RTs and variations from sample preparation and LC-MS runs^17^.

A QTRAP 6500+ (AB Sciex, Concord, ON, Canada) was coupled to an Exion LC AD (AB Sciex, Concord, ON, Canada). MS, MS experiments were done with a Turbo V source (AB Sciex, Concord, ON, Canada) operated with ESI probe. A Phenomenex Luna® amino column (100 mm × 2 mm, 3 μm) was used for separation. The mobile phase A was 1 mM ammonium acetate in chloroform: acetonitrile (1:9), while mobile phase B was 1 mM ammonium acetate in acetonitrile: water (1:1). The gradient started at 20 % MPB and 1% MPC. The MPB progressed from 20 % to 85 % between 0.75 min and 14 min while the MPC ascended from 1 % to 15 % between 11 min to 14 min after which conditions were kept for 0.3 min and then the column was re-equilibrated at initial conditions until 16 min. Two injections were made to accommodate all the MRM transitions of the targeted lipid features. The injection volume was 5 μL for the first acquisition run and 1 μL for the second acquisition run. The column temperature was kept at 35 °C. The injector needle was washed with isopropanol:water:dichloromethane (94:5:1, v, v, v) after each injection^17^. Assigned MRM peaks from the acquired data were integrated using SCIEX OS (version 2.1.6) Software and signals were corrected using proper internal standards.

Blank effects for each analyte were checked by comparing proc blank samples to quality control (QC) samples. The precision and reproducibility of the analytical process were checked using the relative standard deviations (RSDs) of the QCs. With acquisition method 1, 305 analytes with QC RSDs <30% and blank effects < 40% in both QC pools were reported. 58 of the analytes had QC RSDs <15% in both QC pools. With acquisition method 2, 173 analytes with QC RSDs <30% and blank effects < 40% in both QC pools were reported. 70 of the analytes had QC RSDs <15% in both QC pools. With acquisition method 3, 5 TGs with QC RSDs <30% and blank effects < 40% in both QC pools were reported. All data is reported as relative response ratios.

### Reverse phase (RP) lipids platform (ceramides, DAGs, CE)

The RP MS, MS-based lipids platform covers 186 lipids, including ceramides, diglycerides, and cholesterol esters. The identification of metabolites in this platform are based on the dimensions of specific MS, MS transitions and retention time (RT), Using reversed phase columns, the lipids from the same class elute at RTs that can fits into linear regression models involving carbon number and double bond number (RT mapping)^18^. For each class, one or more deuterated internal standards were used to check the RTs and variations from sample preparation and LC-MS runs.

A QTRAP 7500 (AB Sciex, Concord, ON, Canada) coupled to an Exion LC AD (AB Sciex, Concord, ON, Canada). MS, MS experiments were done with a Turbo V source (AB Sciex, Concord, ON, Canada) operated with ESI probe. An Acquity UPLC BEH C8 column (Waters) was used to measure the samples. The gradient was the following: starting conditions 10% B and 10% C; increase of B from 10% to 40% between 1 min and 2 min; maintaining B at 40% and C at 10% between 2 min and 7 min; increase of C from 10% to 45% between 7 min and 8 min; maintaining B at 40% and C at 45% between 8 min and 10 min; returning to initial conditions at 10.5 min and re-equilibration for 1.5 min. The triple quadrupole mass spectrometer operated in polarity switching mode and all analytes were monitored in dMRM mode. Data was acquired using Sciex OS Software V2.0.0.45330 (AB Sciex).

Assigned MRM peaks from the acquired data were integrated using SCIEX OS (version 2.1.6) Software and signals were corrected using proper internal standards.

Blank effects for each analyte were checked by comparing proc blank samples to quality control (QC) samples. The precision and reproducibility of the analytical process were checked using the relative standard deviations (RSDs) of the QCs.

In total 89 analytes with QC RSDs <30% in both QC pools and blank effects < 40% were reported. 43 of these analytes had QC RSDs <15% in both QC pools. The data is reported as relative response ratios.

### Endocannabinoid synthesis lipid platform (NAPE, lysoNAPE, GPNAE, free fatty acids)

The endocannabinoid synthesis platform covers *N*-acyl-phosphatidylethanolamines (NAPEs), 2-lyso-*N*-acyl-phosphatidylethanolamines (lyso-NAPEs), glycerol-phospho-acylethanolamines (GP-NAEs), free fatty acids (FFAs), which profiles the metabolic pathways of *N*-acyl ethanolamines (NAEs).

A QTRAP 6500+ (AB Sciex, Concord, ON, Canada) coupled to an Exion LC AD (AB Sciex, Concord, ON, Canada). MS, MS experiments were done with a Turbo V source (AB Sciex, Concord, ON, Canada) operated with ESI probe. The separation was performed in a BEH C8 column (50 mm × 2.1 mm, 1.7 μm) from Waters Technologies (Mildford, MA, USA) maintained at 40°C, with the flow rate at 0.4 mL, min. The mobile phase was consisted of 2 mM HCOONH_4_, 10 mM formic acid in water (A), ACN (B), IPA (C). The gradient was the following: starting conditions 20% B and 20% C; increase of B from 20% to 40% between 1 min and 2 min; maintaining B at 40% and C at 20% between 2 min and 7 min; increase of C from 20% to 50% between 7 min and 8 min; maintaining B at 40% and C at 50% between 8 min and 10 min; returning to initial conditions at 10.5 min and re-equilibration for 1.5 min. The triple quadrupole mass spectrometer operated in polarity switching mode and all analytes were monitored in dMRM mode. Data was acquired using Sciex OS Software V2.0.0.45330 (AB Sciex).

Assigned MRM peaks from the acquired data were integrated using SCIEX OS (version 2.1.6) Software and signals were corrected using proper internal standards.

Blank effects for each analyte were checked by comparing proc blank samples to quality control (QC) samples. The precision and reproducibility of the analytical process were checked using the relative standard deviations (RSDs) of the QCs.

In total 68 analytes with QC RSDs <30% in both QC pools and blank effects < 40% were reported. 31 of these analytes had QC RSDs <15% in both QC pools. The data is reported as relative response ratios.

### Cholesterol synthesis and oxysterols platform

The cholesterol synthesis and oxysterols platform covers precursors of cholesterol from the Bloch and Kandutsch-Russell pathways, including lathosterol and desmosterol. Oxysterols, which are cholesterol metabolites that can be produced through enzymatic or radical oxidation, are also covered. Reference standards or corresponding deuterated internal standards were used for each analyte for peak identification, 5 deuterated internal standards were used for the correction of variations from sample preparation and LC-MS runs.

A QTRAP 7500 (AB Sciex, Concord, ON, Canada) coupled to an Exion LC AD (AB Sciex, Concord, ON, Canada). MS/MS experiments were done with a Turbo V source (AB Sciex, Concord, ON, Canada) operated with ESI probe. The separation was performed in a BEH C18 column (50 mm × 2.1 mm, 1.7 μm) from Waters Technologies (Mildford, MA, USA) maintained at 40°C, with the flow rate at 0.6 mL, min. The mobile phase was consisted of 0.1% acetic acid in water (A), ACN (B), MeOH (C). The gradient was the following: starting conditions 27% B and 27% C; increase of B & C from 27% to 30% between 1 min and 2 min; increase of B & C from 30% to 48% between 1 min and 2 min; maintaining B & C at 48% between 6.5 min and 7 min; increase of B & C from 48% to 50% between 7 min and 10 min; maintaining B & C at 50% between 10 min and 11 min; returning to initial conditions at 11 min and re-equilibration for 1 min. The triple quadrupole mass spectrometer operated in polarity switching mode and all analytes were monitored in dMRM mode. Data was acquired using Sciex OS Software V2.0.0.45330 (AB Sciex).

Assigned MRM peaks from the acquired data were integrated using SCIEX OS (version 2.1.6) Software and signals were corrected using proper internal standards.

Blank effects for each analyte were checked by comparing proc blank samples to quality control (QC) samples. The precision and reproducibility of the analytical process were checked using the relative standard deviations (RSDs) of the QCs. In total 5 analytes with QC RSDs <30% in both QC pools and blank effects < 40% were reported. Desmosterol and 7-dehydrocholesterol had high RSDs in lesion QC pools, but acceptable RSDs in white matter QC pools and were kept. The data is reported as relative response ratios.

### Oxidized phospholipids platform

The oxidized phospholipids (OxPLs) platform covers a class of lipid molecules formed when phospholipids undergo oxidative modification. Reference standards of oxidized phosphatidylethanolamines (oxPCs) were used for peak identification. More OxPCs and OxPEs were identified based on MS, MS transitions and HRMS, MS at positive mode. Deuterated internal standards of PCs, LPCs and eCBs were used for the correction of variations from sample preparation and LC-MS runs.

A QTRAP 7500 (AB Sciex, Concord, ON, Canada) coupled to an Exion LC AD (AB Sciex, Concord, ON, Canada). MS, MS experiments were done with a Turbo V source (AB Sciex, Concord, ON, Canada) operated with ESI probe. The separation was performed in a BEH C8 column (50 mm × 2.1 mm, 1.7 μm) from Waters Technologies (Mildford, MA, USA) maintained at 40°C, with the flow rate at 0.4 mL, min. The mobile phase was consisted of 2 mM ammonium acetate, ACN (B), IPA (C). The gradient was the following: starting conditions 10% B and 10% C; increase of B from 10% to 45% between 1 min and 2.5 min; maintaining B at 45% and C at 10% between 2.5 min and 4 min; increase of C from 10% to 48% between 4 min and 7.5 min; maintaining B at 45% and C at 48% between 7.5 min and 10 min; returning to initial conditions at 10.5 min and re-equilibration for 1.5 min. The triple quadrupole mass spectrometer operated in polarity switching mode and all analytes were monitored in dMRM mode. Data was acquired using Sciex OS Software V2.0.0.45330 (AB Sciex).

Assigned MRM peaks from the acquired data were integrated using SCIEX OS (version 2.1.6) Software and signals were corrected using proper internal standards.

Blank effects for each analyte were checked by comparing proc blank samples to quality control (QC) samples. The precision and reproducibility of the analytical process were checked using the relative standard deviations (RSDs) of the QCs.

In total 26 analytes with QC RSDs <30% in both QC pools and blank effects < 40% were reported. The data is reported as relative response ratios.

To estimate absolute concentrations, a calibration line containing OxPCs and PC (16:0, 20:4) was prepared separately from the study samples. A response (area ratio) – concentration relationship was established with the calibration line, and the concentrations of OxPCs and PC (16:0, 20:4) were calculated, but only for those lipids for which a standard was available.

### Chemical proteomics (ABPP) sample preparation

The chemical proteomics workflow was based on the previously reported procedures by Van Rooden *et al.*^84^ Lysates were prepared by glass-bead mechanical lysis of isolated lesions in 20 mM HEPES, 1 mM MgCl_2_, 2 U/mL benzonase. After protein quantification using the Bradford method (Bio-Rad), lysates were snap-frozen in LN_2_ and stored at -80 ⁰C until further use.

Lysates (100 µL, 1mg/mL protein) were thawed on ice and subsequently treated with a cocktail of 10 µM FP-biotin and 10 µM THL-biotin as described previously.^84^ As controls, 10 heat and SDS inactivated (1% SDS, 5 minutes 95 ⁰C) lysates were taken along in the procedure. Probe treated lysates were precipitated using the MeOH, CHCl_3_ precipitation. The protein pellet was washed in MeOH and subsequently redissolved in PBS containing 0.5% SDS and 5 mM DTT. Proteins were dissolved by sonication and the solution was incubated at 65 ⁰C for 15 min to reduce cysteines. Afterwards, cysteines were alkylated using iodoacetamide and excess iodoacetamide was quenched using DTT. Subsequently, the samples were added to a suspension of 5 µL high-capacity streptavidin agarose beads (Thermo-Fisher, 20361) and 15 µL control agarose beads in PBS (Thermo Fisher, 26150) containing 0.25% SDS. The suspension was incubated for 2 hours at RT while rotating head-over-head to let probes proteins bind to the beads. After incubation, the beads were washed with PBS containing 0.5% SDS (4x) and subsequently PBS (5x). On-bead digestion was performed with 0.25 µg sequence-grade trypsin (Promega) in 100 mM Tris, 100 mM NaCl, 1 mM CaCl_2_ and 2% (vol/vol) acetonitrile) overnight at 37 ⁰C while shaking at 1000 rpm. Trypsin was quenched with 10 µL formic acid. Peptides were then desalted on Oasis plates (Waters) and eluted in 60% acetonitrile with 0.5% formic acid. The solvent was evaporated in a SpeedVac (45 ⁰C, Eppendorf) and samples were stored at -80 ⁰C until further use.

### LC-MS/MS for chemical proteomics (ABPP)

Samples were randomized and assigned to six blocks of 23 samples per block. Each block also contained 4 reference samples that were measured in every measurement-block as described by Zhang *et al.*^85^ Per block of samples, peptides were then reconstituted in 3% acetonitrile in H_2_O and analyzed on a QExactive LC-MS/MS (ThermoFisher). All gradients and solutions were according to previously published procedures.^84^ Before and after each block, the LC/MS machine was cleaned, recalibrated and its performance was assessed.

### Data analysis ABPP with chemical proteomics

Raw spectral data was analyzed by MaxQuant software developed by Cox and Mann using largely default settings.^86^ Match between runs was enabled. The ‘proteingroups.txt’ output file from MaxQuant was imported in R. Identified proteins were filtered for potential contaminants identified by MaxQuant, and additional criteria for the proteins were that 1) a protein was identified with 2 independent unique peptides, 2) The ratio of protein raw intensity of native, heat inactivated lysate was at least 2.0 in at least 5 out of 10 QC pairs. 3) The protein is annotated as a serine hydrolase, or has a annotation in Uniprot as “charge-relay system” or “nucleophile” as a catalytic residue and 4) The protein had an LFQ value for more than 60% of the samples in at least one biological group (Lesion type). 97 enzymes met these criteria and were further processed.

The LFQ values were then corrected for LC/MS performance by block design following the procedure presented by Zhang et al.^85^ In short, the median log2 LFQ for a protein in the reference samples was compared to the same samples in other blocks. The difference in log2 median was substracted from the same protein in the experimental samples. Analysis of the reference samples themselves showed that this correction improved the pearson’s correlations of the reference samples to each other (data not shown). All subsequent analysis was then performed using the corrected log2 transformed LFQ values.

Missing values were imputed using the random gaussian distribution centered around the minimum LFQ value found for that protein in the dataset, with a standard deviation equal to the 1/3 of the standard deviation of that protein in the dataset. This is based on the assumption that if a protein was not found, it was most likely below the detection threshold, and therefore very low.

Enzyme activity values were associated with the MOFA factors derived from the other datasets using Limma. The ABPP dataset was log2 transformed, centered around 0, and then fed into a linear model using the design y ∼ f1 + f2 + f3 + f4 + f5 + f6 + f7. P-values for the derived coefficients were calculated using the eBayes function, and subsequently these p values were adjusted for multiple testing using the Benjamini-Hochberg method with a false discovery rate set at 10%.

