## Supplementary material for "Foamy microglia link oxylipins to disease progression in multiple sclerosis": vlietetal_2024_supplementary figures

# to

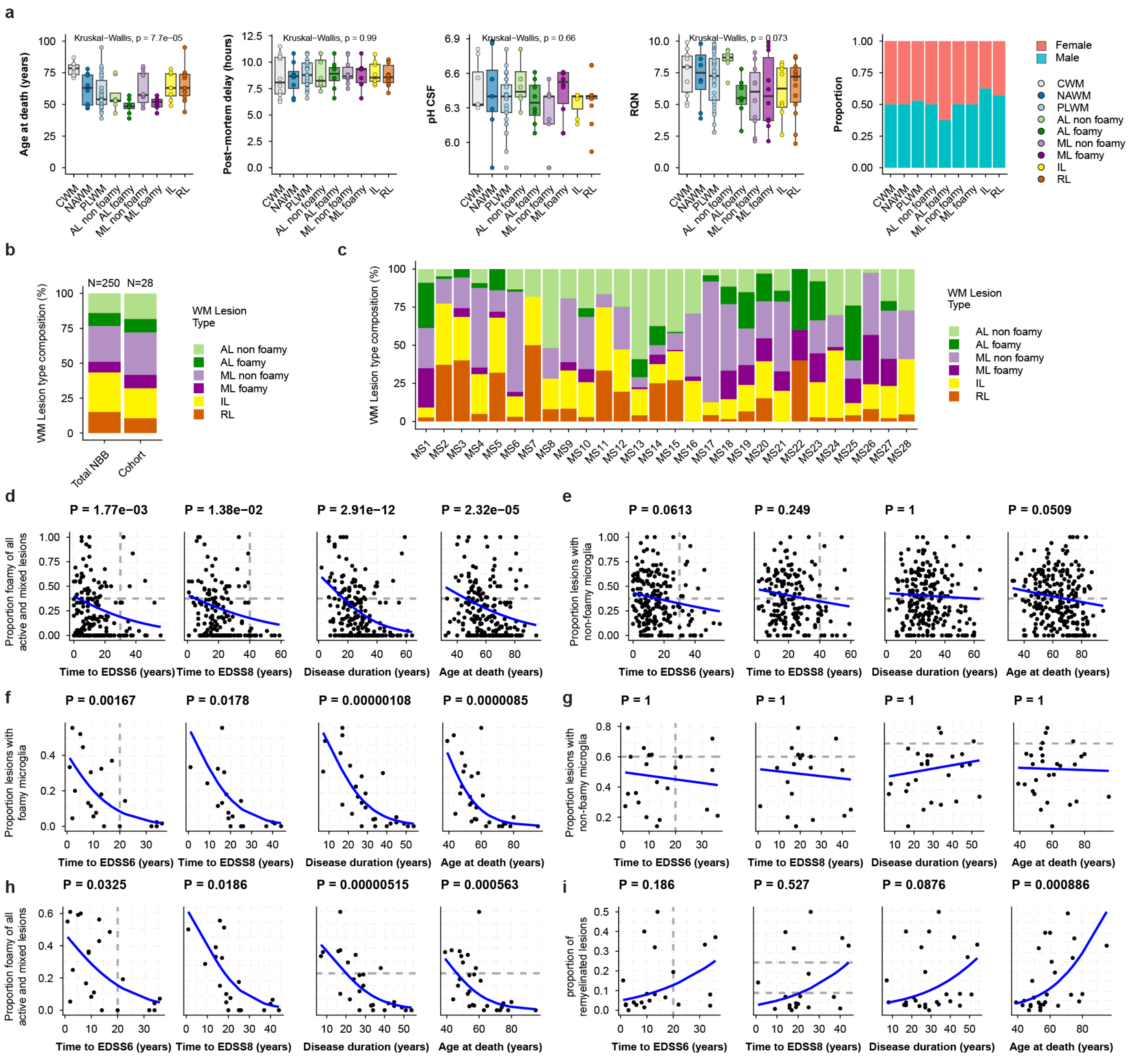


**Fig. S1 Clinical characteristics of the selected cohort. a,** baseline characteristics of the donors. **b,** distribution of lesion types across all lesions in the donors from the entire Netherlands brain bank (N=250) or the cohort analyzed here (N=28). **c,** distribution of lesion types within individual donors shows variability of lesion types between individuals. **d-i,** Association of lesion type proportion with disease progression parameters for the entire NBB cohort (**d-e**) or the cohort analyzed here (**f-i**). A generalized linear model was used to model the association using a quasi-binomial distribution for lesion type proportion, with a likelihood-ratio test to obtain a p value, which was corrected for multiple testing.

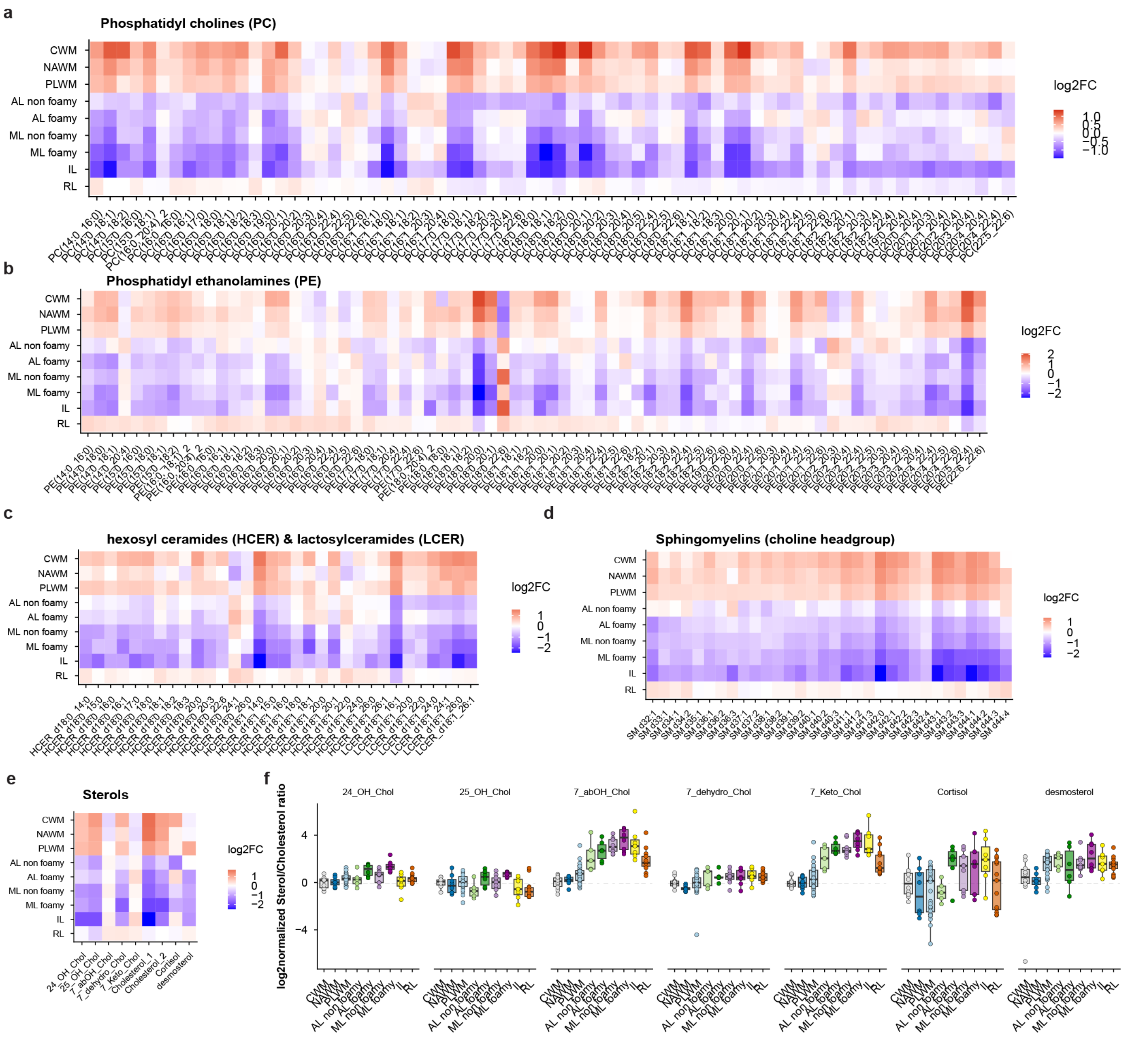


**Fig. S2. Lipidomics shows loss of lipids in MS lesions. a-e** Data represent the normalized mean log2 transformed levels across lesion types for phosphatidyl choline (PC, **a**), phosphatidyl ethanolamine (PE, **b**), cerebrosides HCER and LCER (**c**), sphingomyelins (**d**) and sterols (**e**). **f,** the change in sterol/cholesterol ratios across lesion types. We lack absolute quantification of these lipids, so we can not report the absolute ratio, only the relative change in ratio.

a

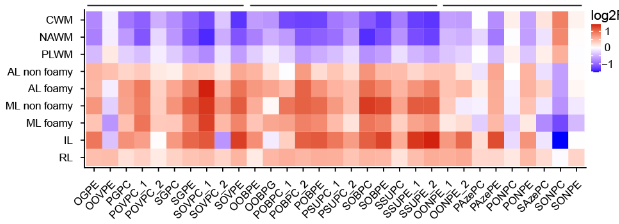


b

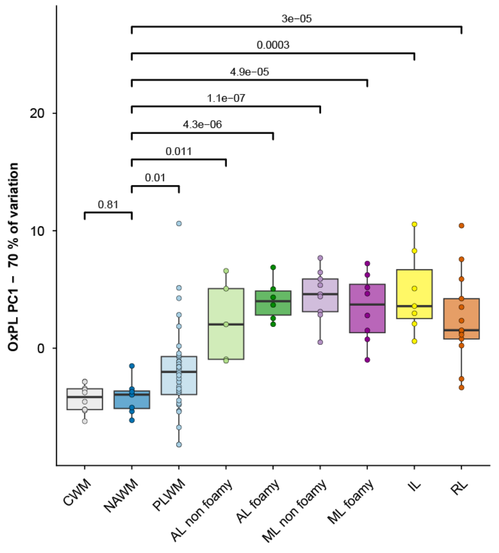


c

absolute concentration oxPCs

| Lipid | pmol/mg | ~conc. (µM) |
|---|---|---|
| POVPC | 1.5 | 1.5 |
| PAzePC | 0.72 | 0.72 |
| POBPC | 0.028 | 0.028 |
| PGPC | 0.14 | 0.14 |
| PSUPC | 0.038 | 0.038 |
| PAPC (PC) | $2.0*10^3$ | $2.0*10^3$ |

d e

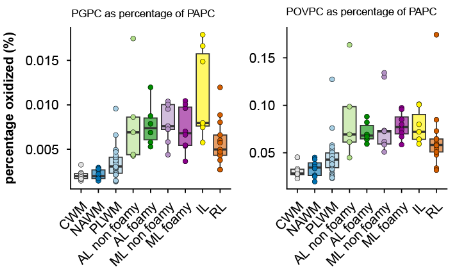


f

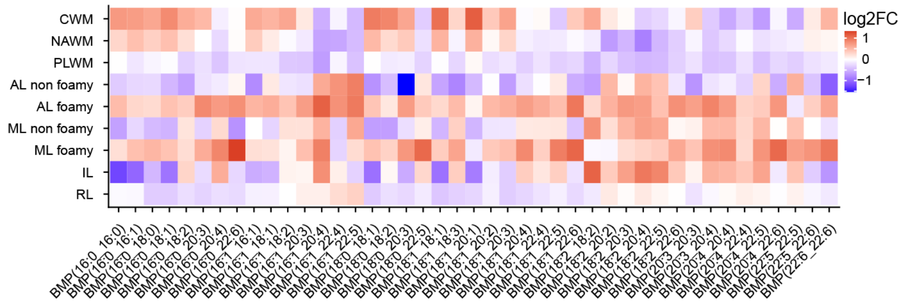


**Fig. S3. Oxidized phospholipid levels and bismonoacylglycerolphosphates. a,** oxidized phosholipid levels across lesions. Data represent the normalized mean log2 transformed levels. **b,** the first principal component of the oxPL levels across lesion types, summarizing all oxPL species into one score. **c,** mean absolute levels of selected oxPL species in MS lesion. The concentration is reported as pmol per mg tissue, which is roughly equivalent to µM assuming a density of 1mg/mL. The originating PC precursor (PAPC) is given as reference. **d-e,** the percentage of PAPC which is oxidized to PGPC (**d**) or POVPC (**e**). **f,** bismonoacylglycerolphosphate (BMP) levels across lesions. Data represent the normalized mean log2 transformed levels.

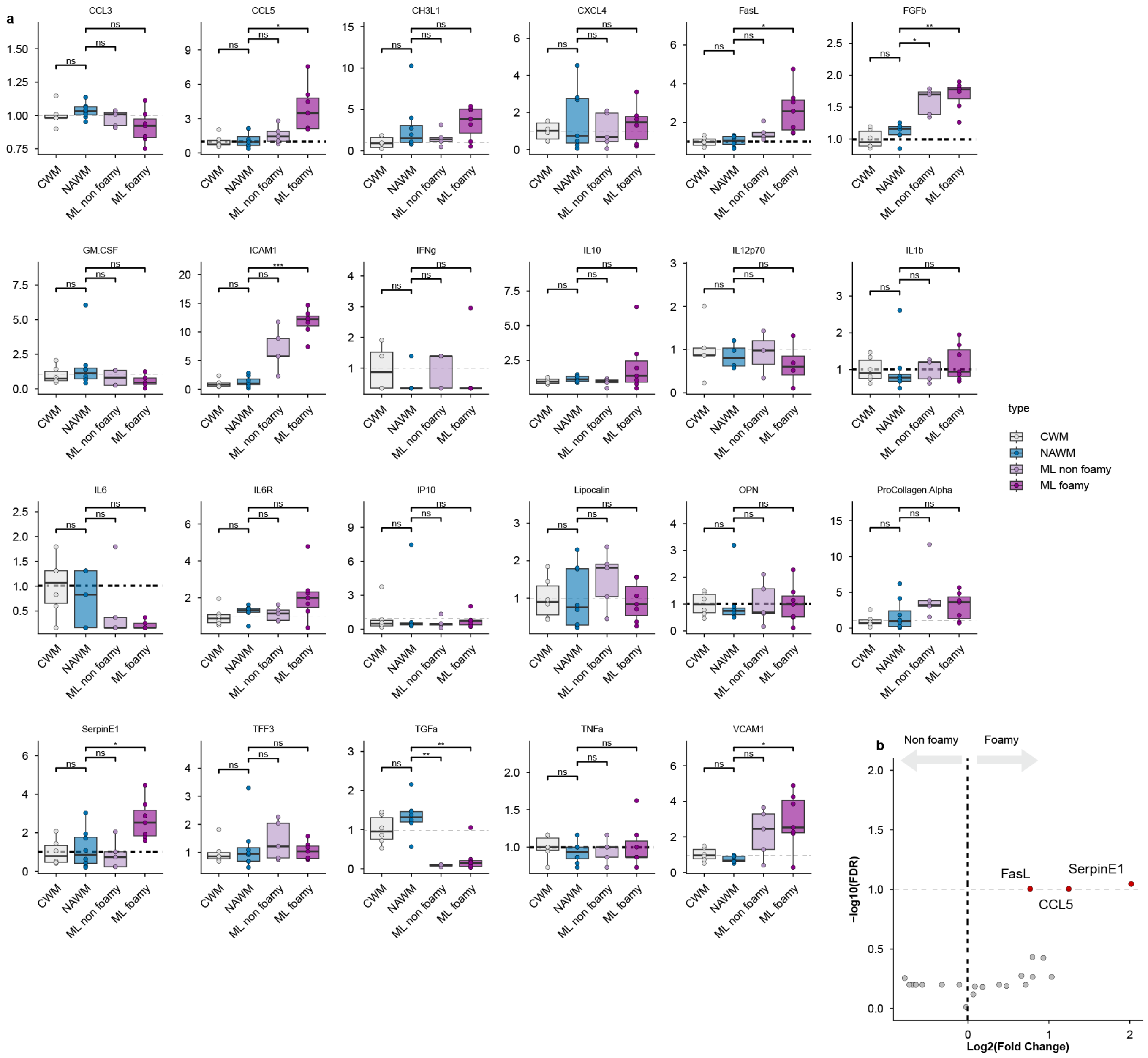


**Fig. S4. Cytokine screening. a,** Boxplots of cytokine concentrations by sample type normalized on the control mean. **b,** volcanoplot of cytokine differences between ML with foamy compared ML with non-foamy microglia. p values were calculated by Limma and BH-corrected for multiple testing

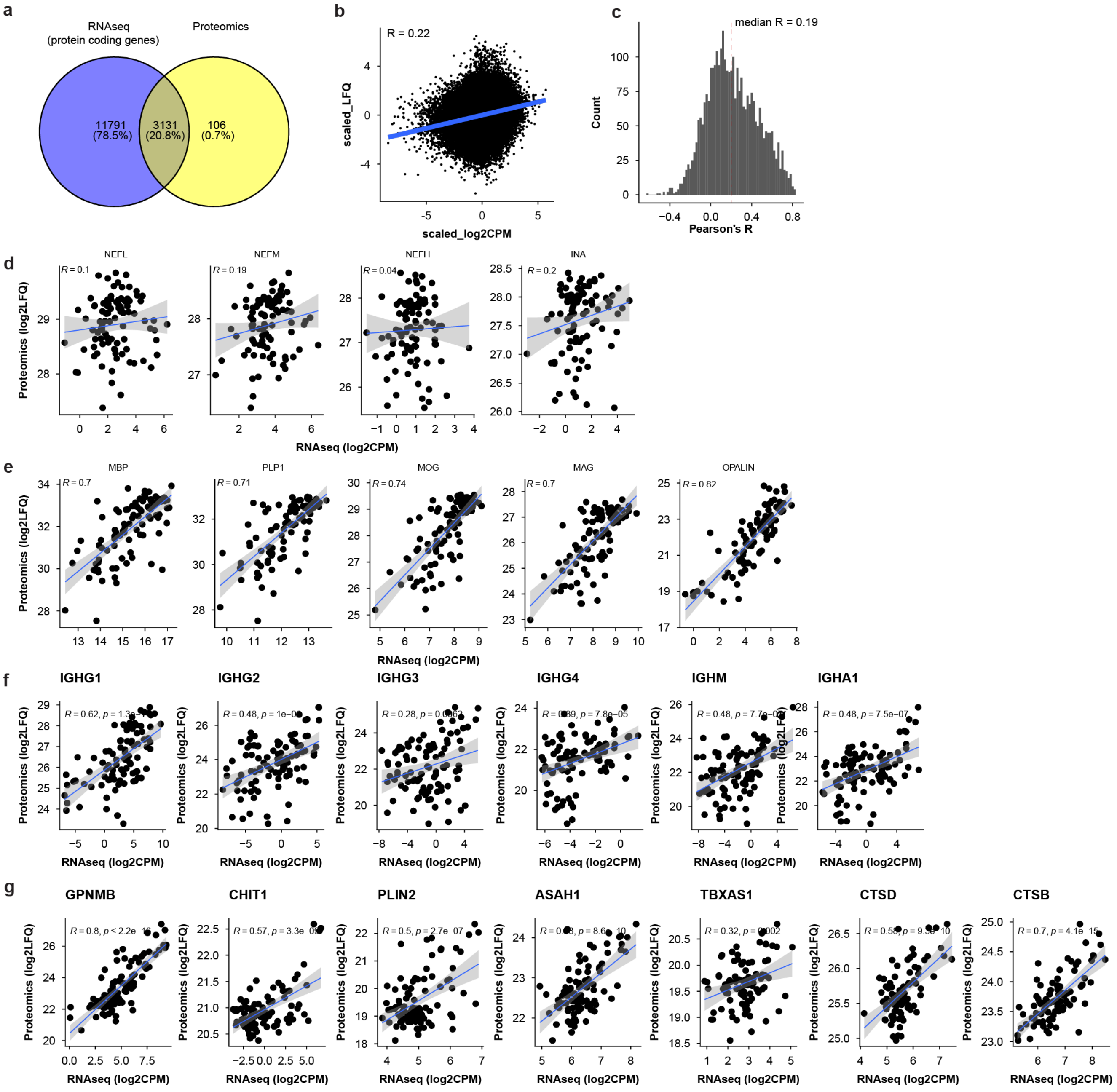


**Fig. S5. RNA protein correlation in MS lesions. a,** overlap between protein-coding genes detected by RNA sequencing or proteomics. **b,** global correlation of RNA and protein levels. All genes/proteins were z-scored and a pearson correlation coeffiicient was calculated on the global level, with all genes/protein added together. **c,** correlations were calculated (pearson) for Individual gene-protein pairs and the obtained R values were plotted in a histogram, showing that some genes have no correlation with their protein product, while other genes are strongly correlated. **d-g,** example correlations of proteins as calculated in **c** for proteins of the axon (**d**), myelin sheeth (**e**), immunoglobulins (**f**) or protein related to foamy microglia (**g**).

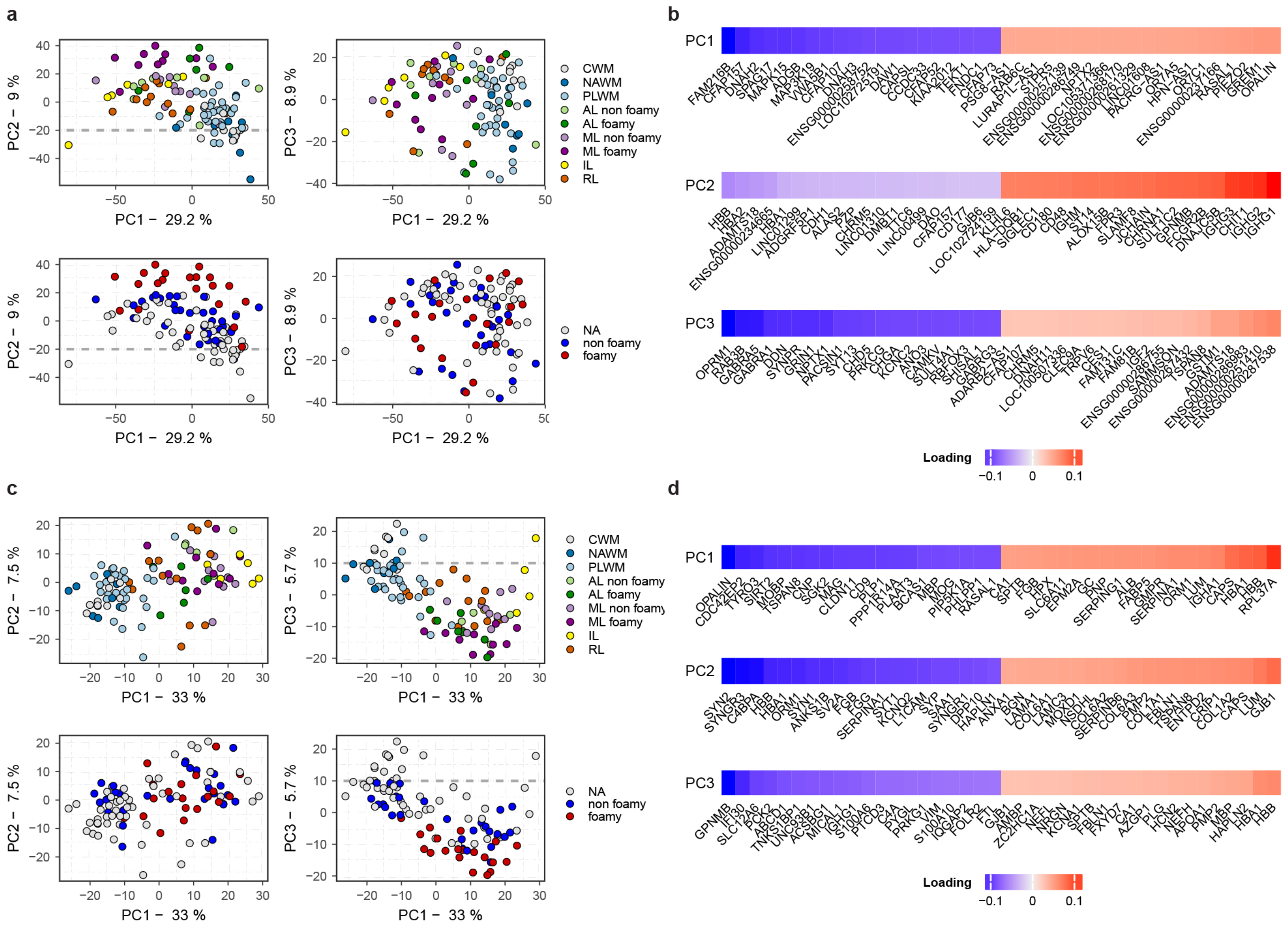


**Fig. S6. PCA analysis of the RNAseq and proteomics data sets. a+c,** PCA plots showing the first 3 principal components for the RNAseq data (**a**) or the proteomics data (**c**), coloured by either lesion type or morphology. **b,** top 20 positive and negative loadings for the first three principal components for the RNAseq data (**b**), or the proteomics data (**d**).

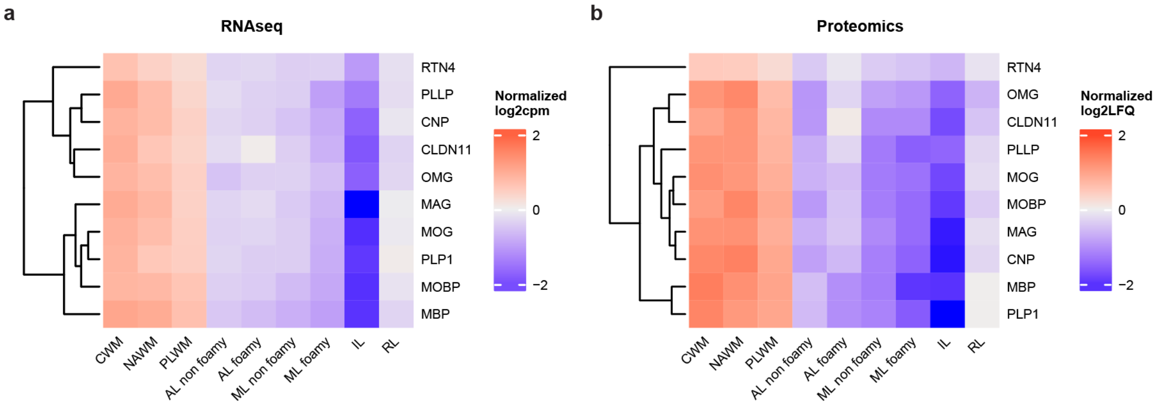


**Fig. S7. Myelin levels decrease in all lesion types. a,** Mean myelin gene expression (**a**) of protein levels (**b**) across lesion types.

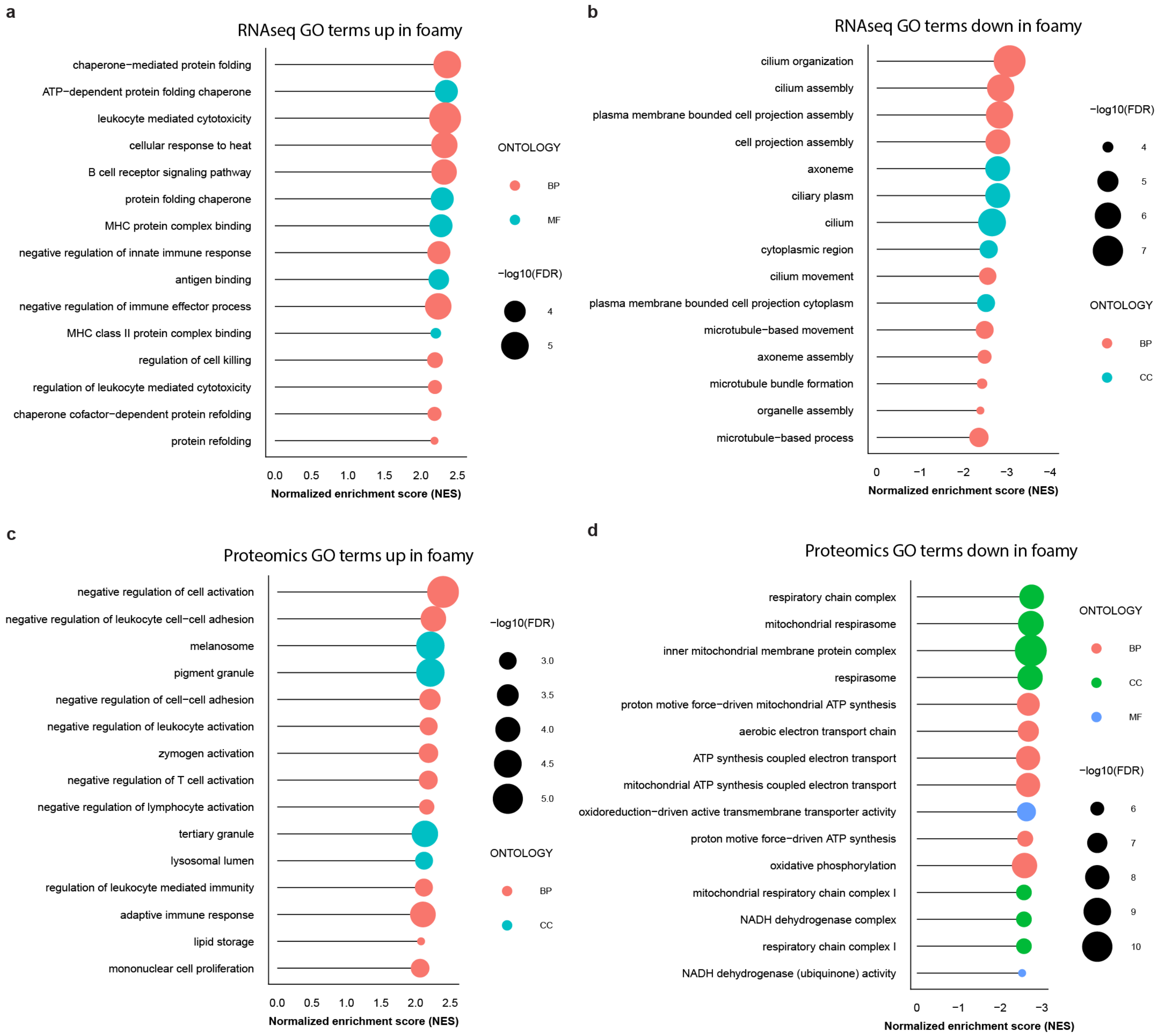


**Fig. S8. Pathways associated with foamy microglia compared to non-foamy microglia.**

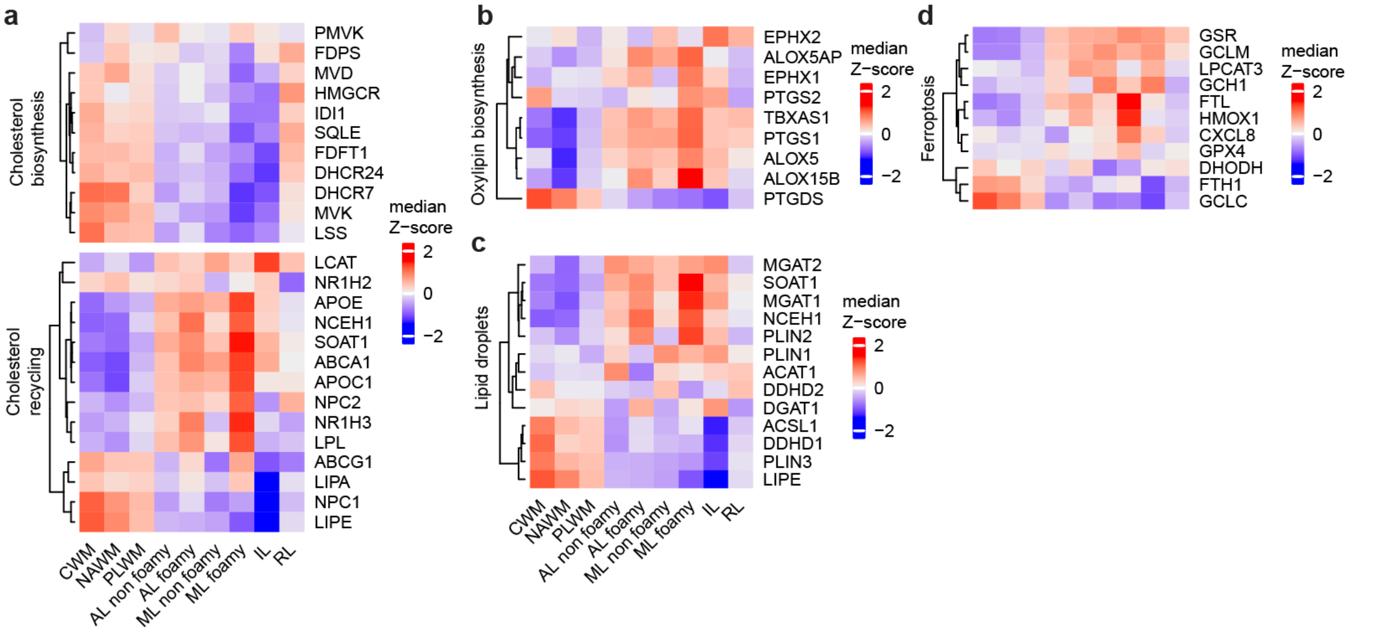


**Fig. S9. Genes involved in lipid metabolism are increased in lesions, most strongly in lesions with foamy microglia.** Heatmaps depicting Z-scores of gene expression of genes involved in cholesterol synthesis and recycling (a), oxylipin biosynthesis (b), lipid droplet formation (c) and ferroptosis and lipid peroxidation (d).

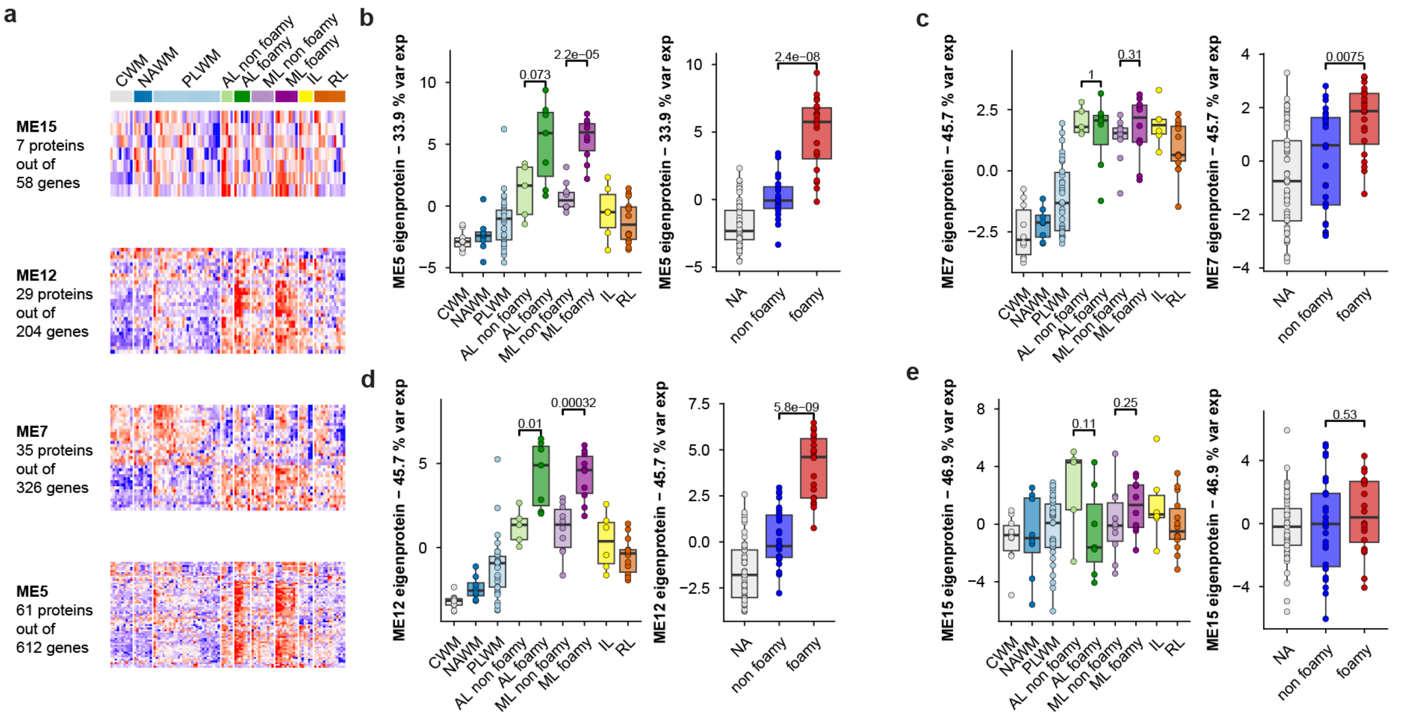


**Fig. S10. Replication of WGCNA modules on the protein level. a,** Genes from the four modules associated with ML with foamy (Figure 3e) microglia were extracted from the WGCNA, and their protein products were searched in the proteomics dataset. A clustered heatmap depicting Z-scores of those proteins per module is depicted. **b-e,** analogously to the eigengene method in WGCNA, an “eigenprotein” was calculated for the proteins corresponding to one gene-module and the first principal component is plotted in a boxplot, with a Wilcoxon rank sum test to assess significance between lesion types or morphology.

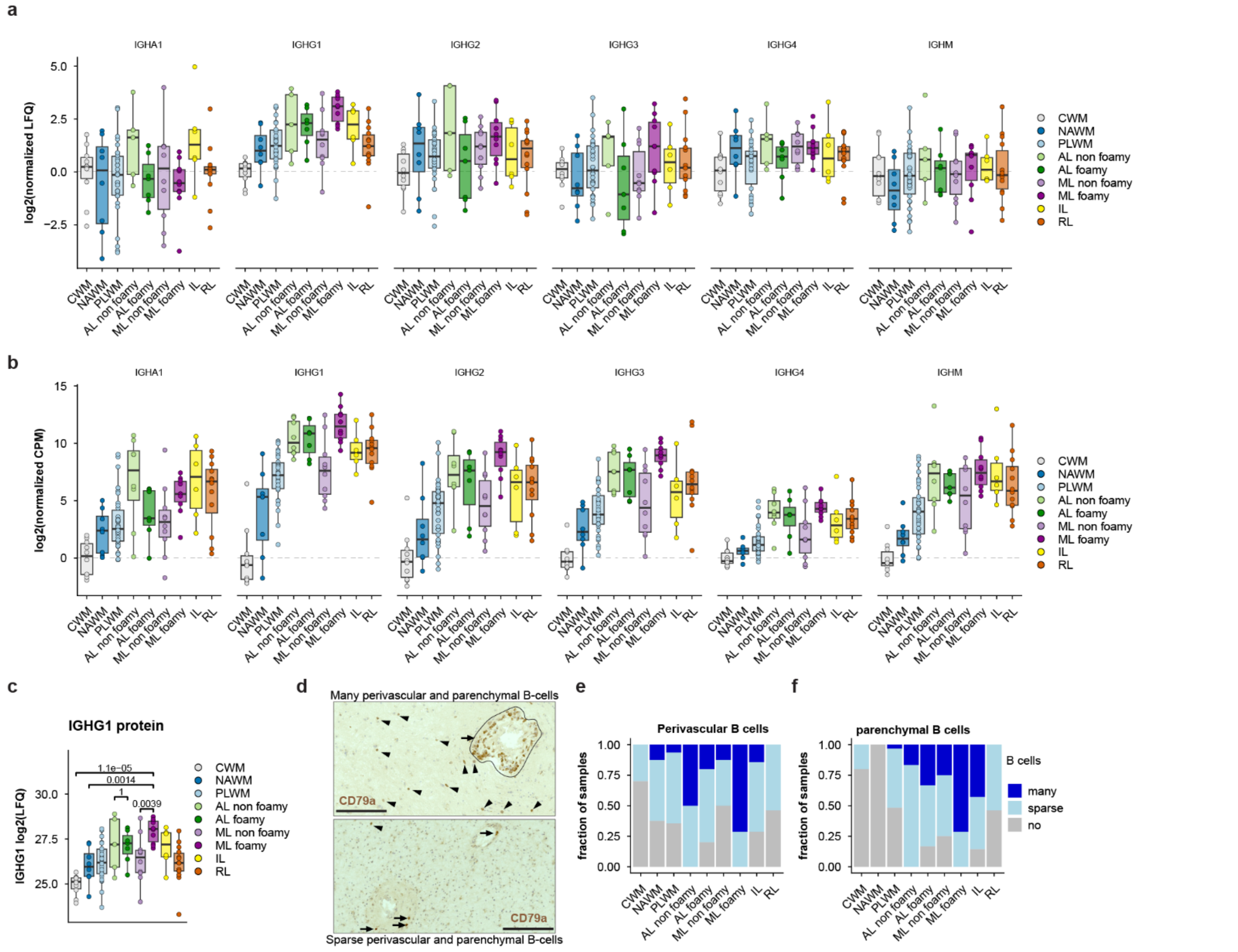


**Fig. S11. B-cells and immunoglobulin expression across lesion types. a,** Levels of immunoglobulin constant regions as determined by proteomics (**a**) or RNA sequencing (**b**). **c,** IgG1 (*IGHG1*) is strongly upregulated in lesions compared to CWM and NAWM, especially high in ML with foamy microglia. IGHG1 protein levels are also significantly higher in ML with foamy microglia compared ML with non-foamy microglia, this is not the case for active lesions. **d,** representative images of MS lesion with high perivascular and infiltrating B-cells (top) and sparse B-cells (bottom). Arrowheads indicate parenchymal cells, while arrows indicate perivascular cells **e-f**, Lesions were scored as having many, sparse or no B-cells in the perivascular space (**e**) or infiltrating into the parenchyma (**f**).

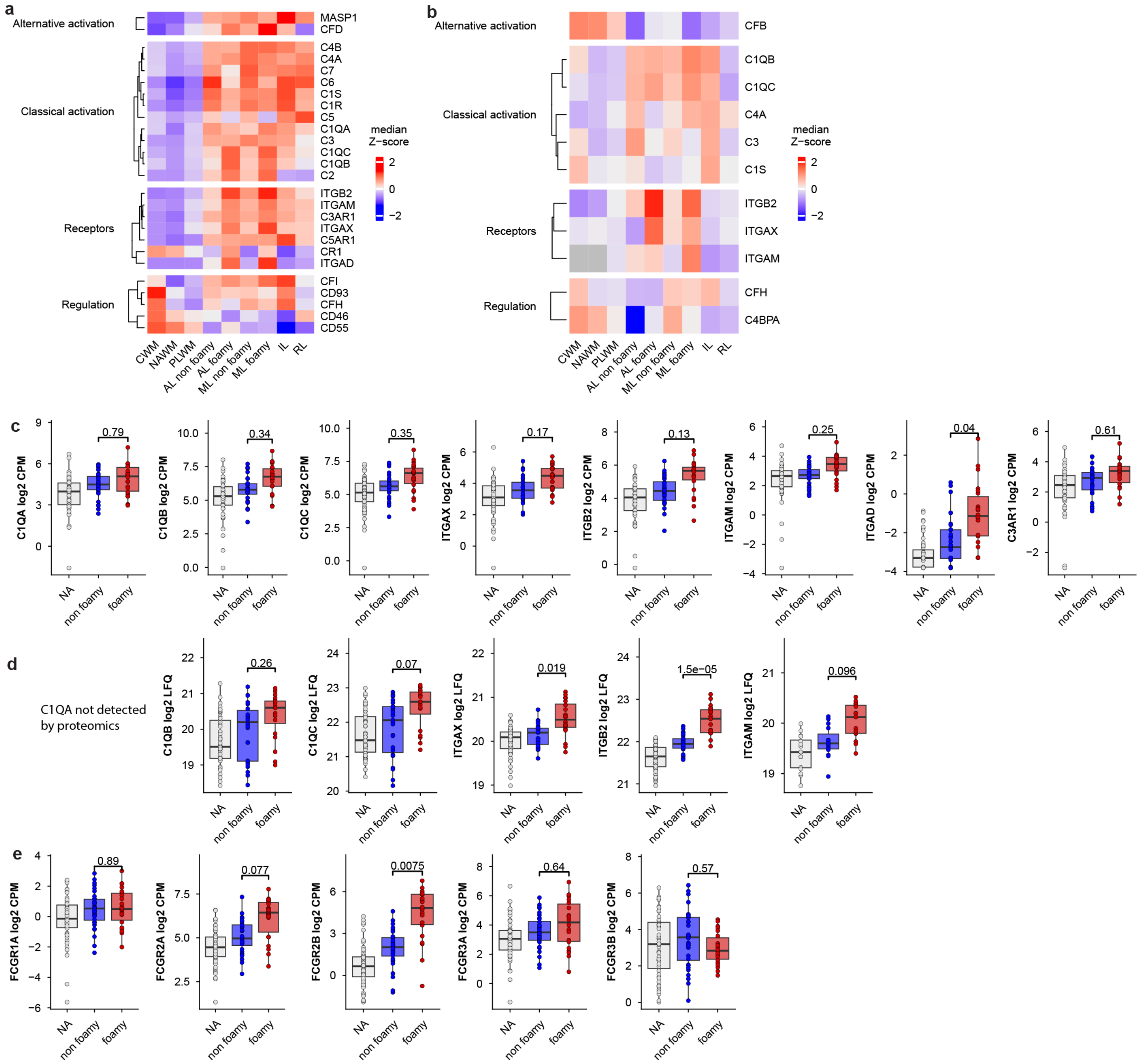


**Fig. S12. Expression of antibody-mediated phagocytosis pathways. a-b,** Heatmap depicting genes (**a**) or proteins (**b**) of the complement system. Data is expressed as the median Z-scored log2 counts per million (**a**) or LFQ (**b**) per lesion type. **c-d,** expression of C1Q genes , as well as iC3b receptors on the gene level (**c**) or protein level (**d**). **e**, Gene expression of Fc gamma receptors. Statistics are calculated using limma with BH-correction for multiple testing (FDR<10%).

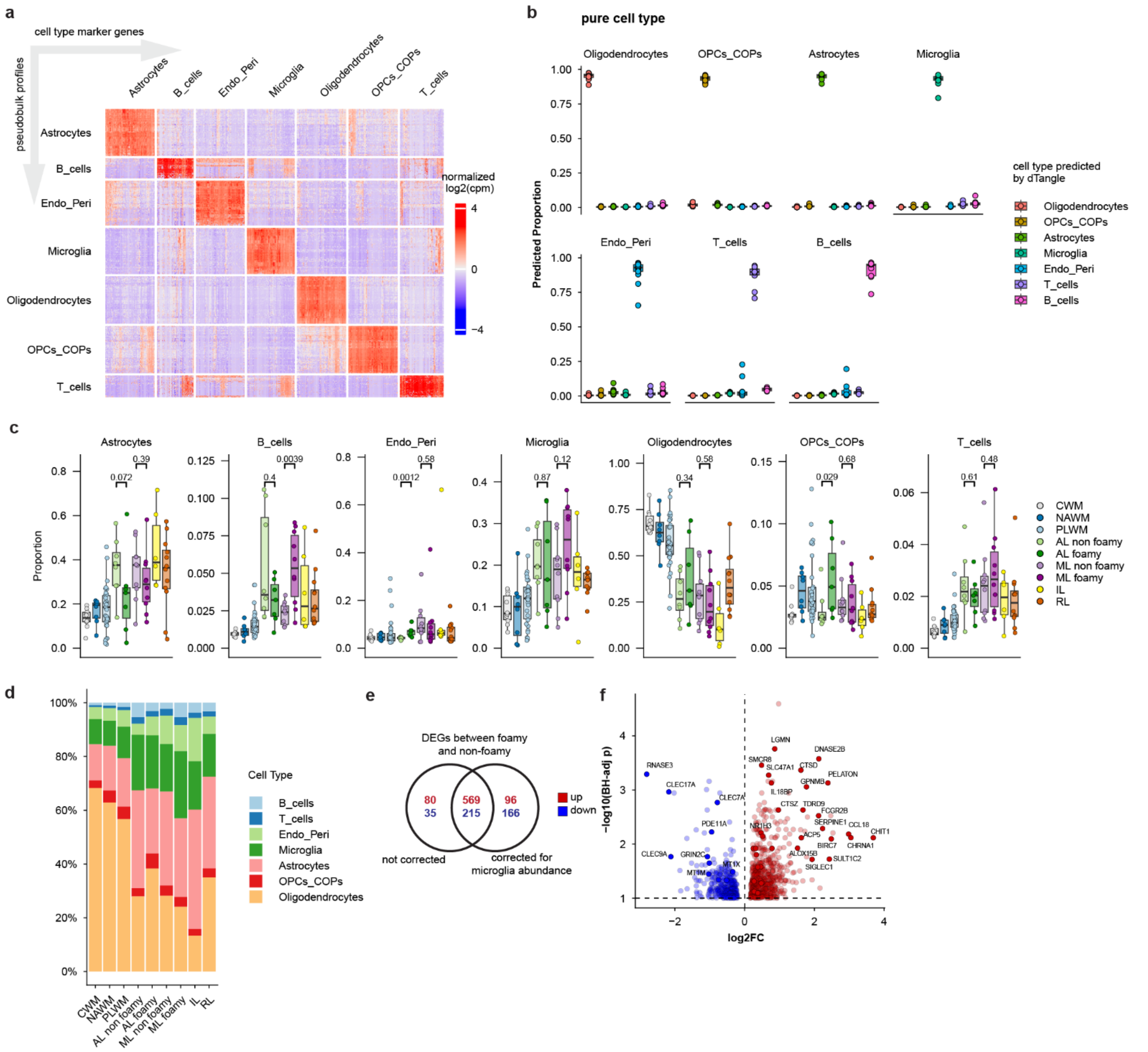


**Fig. S13. Single cell deconvolution of the bulk RNAseq data using snRNAseq. a,** Marker genes selected for each cell type for use in deconvolution. **b,** validation of the method: the snRNAseq was randomly split 80/20, and subsequently 80% of the data was used to predict the cell type composition of the remaining 20% of samples. For each cell type we consistently see a very high percentage (~90%), validating the method. **c,** Boxplots showing the fraction of each cell type across the different lesion types. Statistical test is the wilcoxon rank sum test without any correction for multiple testing. **d,** Global cell type composition of each lesion type. **e,** overlap between differentially expressed genes between foamy and non-foamy samples with and without correction for the microglia numbers in d. **f**, volcanoplot showing DEGs between foamy and non-foamy samples with correction for the microglia numbers in **c.**

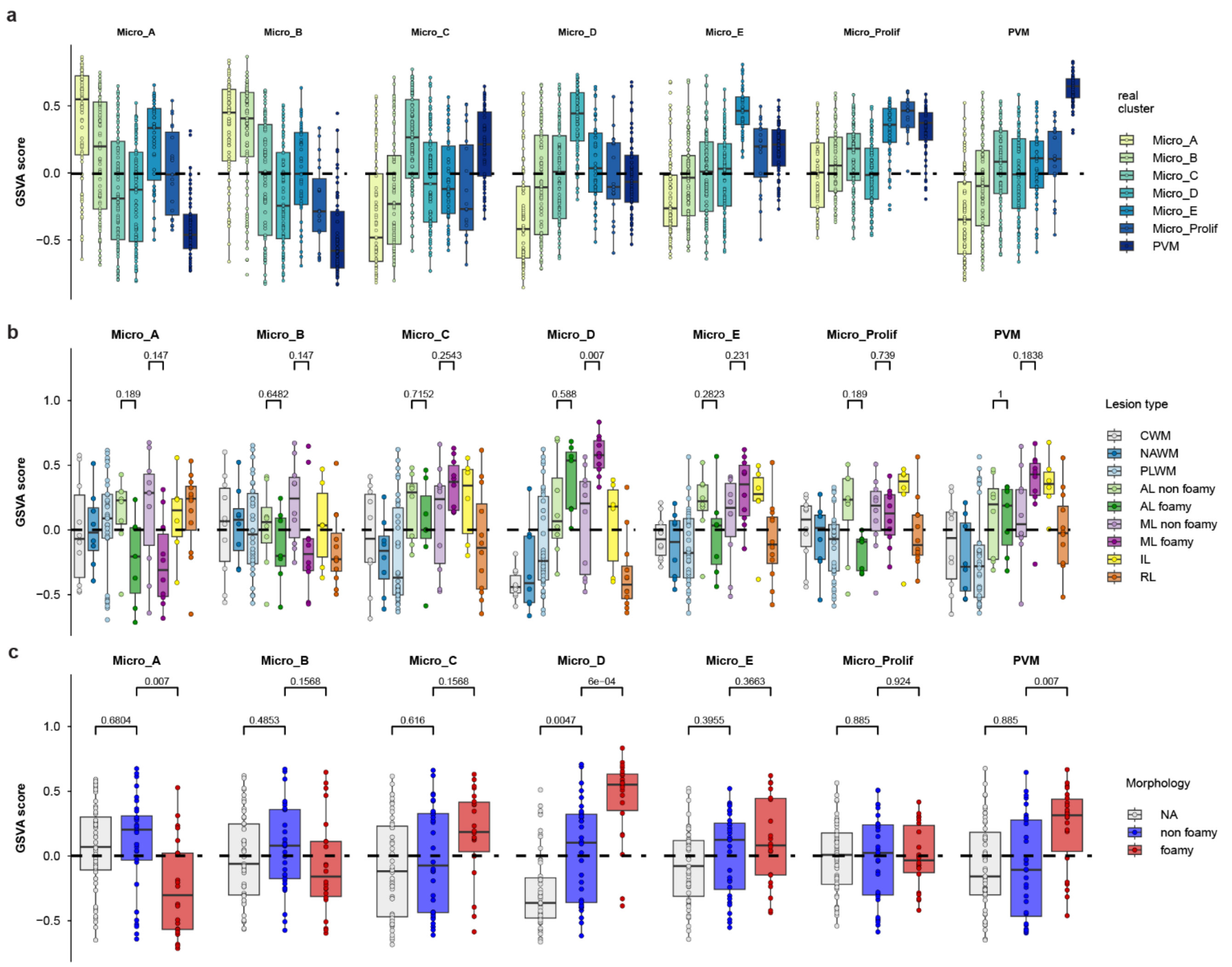


**Fig. S14. The Micro_D state is enriched in foamy MS lesions. a,** validation of the method. Marker genes for each microglial state enriched their own microglial state, except for micro_B. **b-c,** boxplots showing enrichment of each state using GSVA. Statistical test is a wilcoxon rank sum test with BH correction for multiple testing.

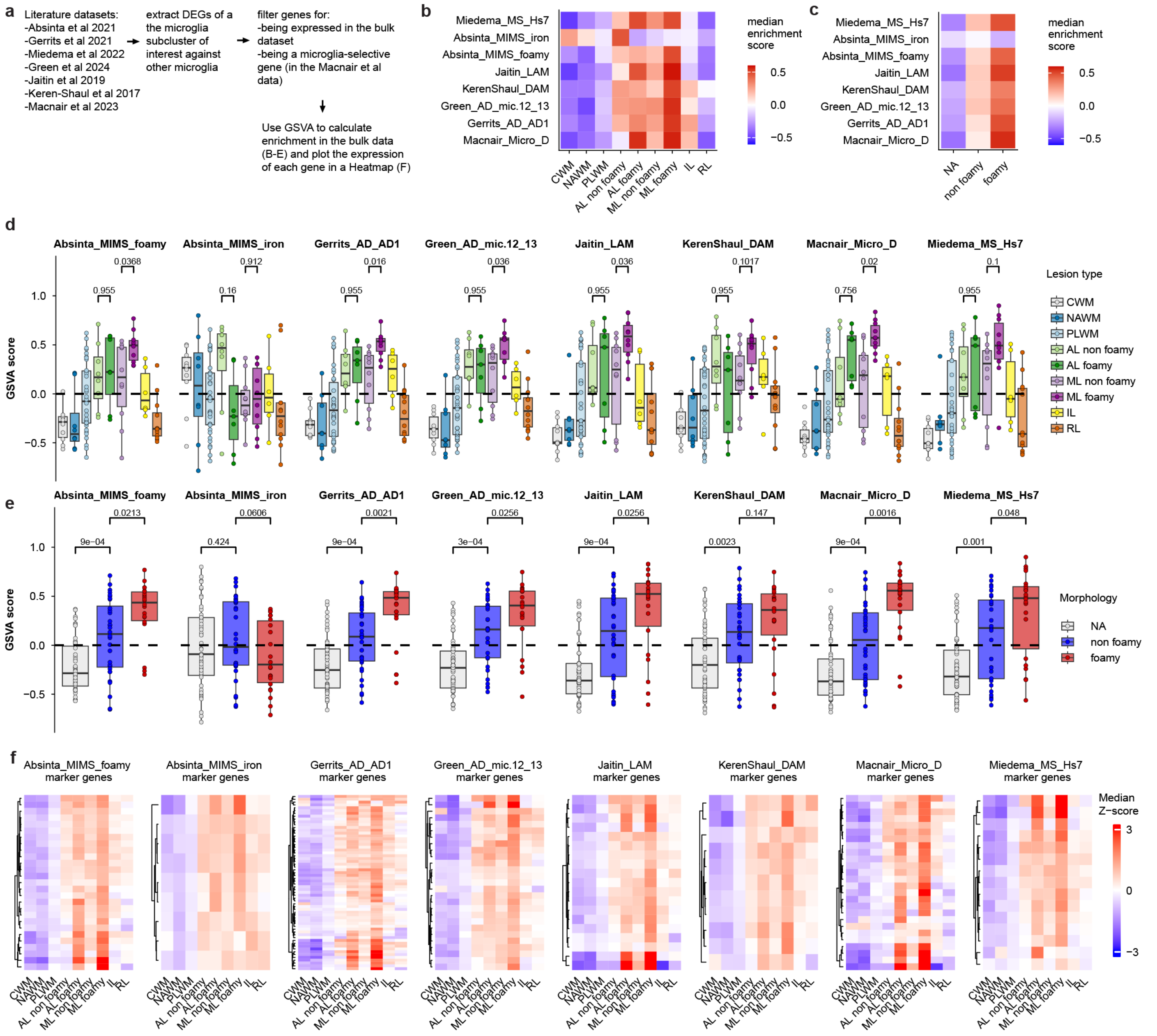


**Fig. S15. Literature microglia states associated with MS, AD and obesity are enriched in foamy MS lesions. a,** Marker genes selection strategy. **b-c,** Heatmaps showing the median enrichment score for each MS lesion type (b) or morphology (c) using GSVA. **d-e,** boxplots showing enrichment of each state using GSVA. Statistical test is the wilcoxon rank sum test with BH correction for multiple testing. **f,** median expression of state-marker genes in the different lesion types. Data is presented as z-scores log2 counts per million.

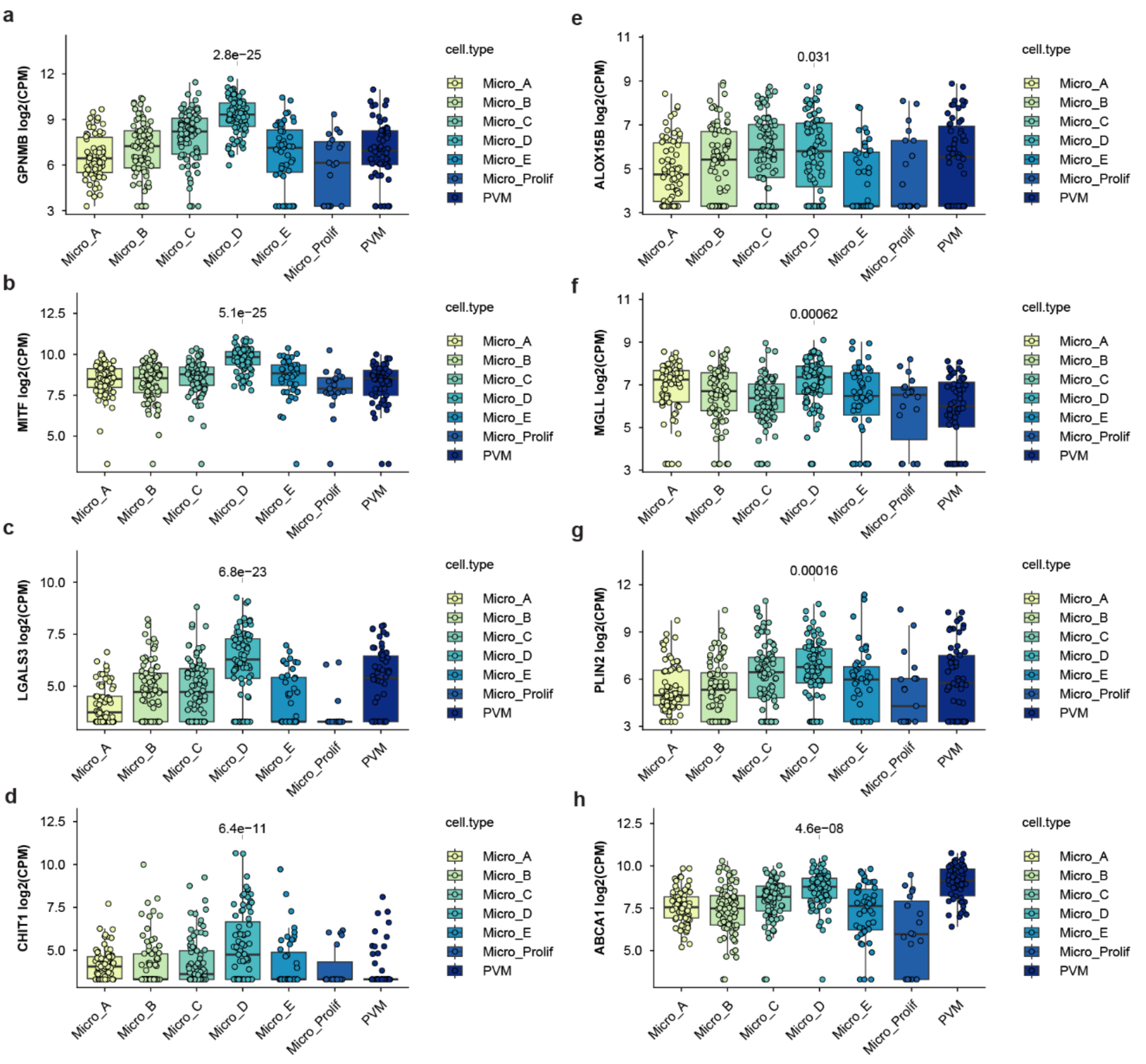


**Fig. S16. Marker genes for the Micro_D microglia state. a-d,** top marker genes for Micro_D. **e-h,** selected genes involved in lipid metabolism that are enriched in the Micro_D profile. P-values represents the comparison of Micro_D to the combined other profiles. P-values were calculated using Limma and BH-corrected for multiple testing.

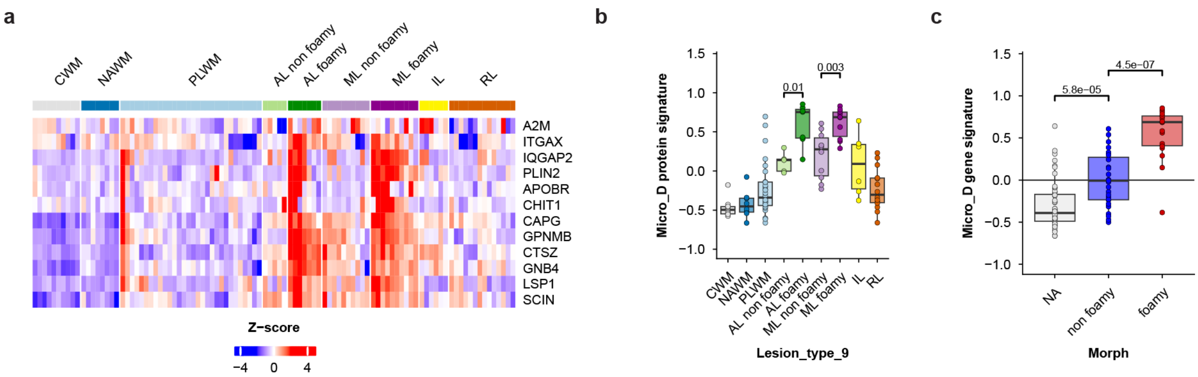


**Fig. S17. Micro_D marker genes on the protein level. a,** Protein Z-scores of proteins encoded by Micro_D marker genes shows strong upregulation in lesions with foamy microglia. **b-c,** The proteins from **a** were summarized into one protein signature score using gene set variation analysis (GSVA). Statistics represent p-values from Wilcoxon rank sum tests.

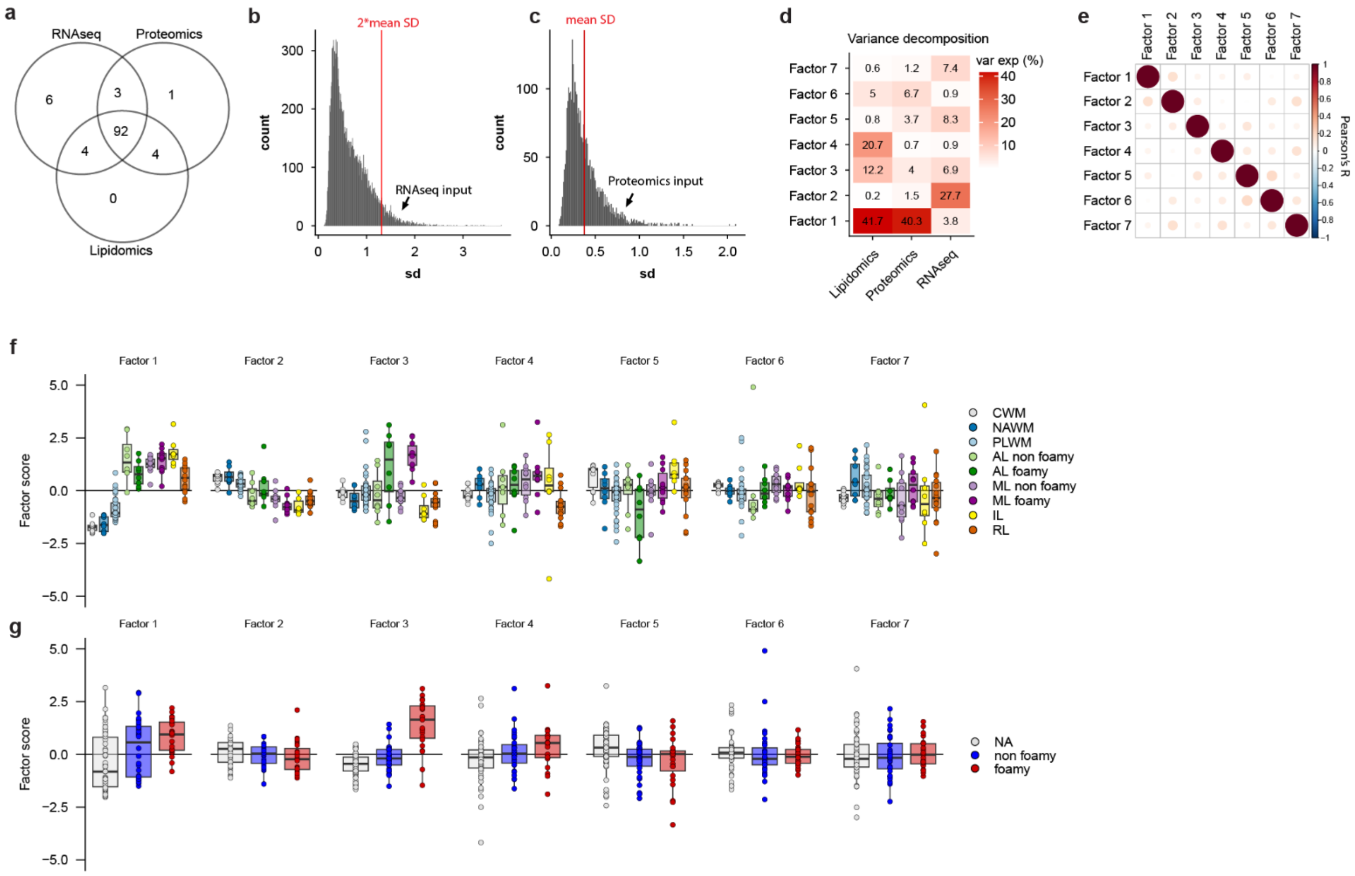


**Fig. S18. Multi-omics data integration. a,** overview of the sample-overlap between the different data modalities. Most samples have been analyzed by all three omics-technologies. However, for some samples there was not enough material, or the sample was discarded because of quality control. **b-c,** data input for RNAseq and proteomics was restricted to highly variable genes (**b**) or proteins (**c**). **d,** variance decomposition of the seven derived MOFA factors. **e,** Pearson correlation of the MOFA factors shows factors are mostly orthogonal to each other. **f-g,** boxplots showing the distribution of MOFA factor scores across lesion type (**f**) or morphology (**g**).

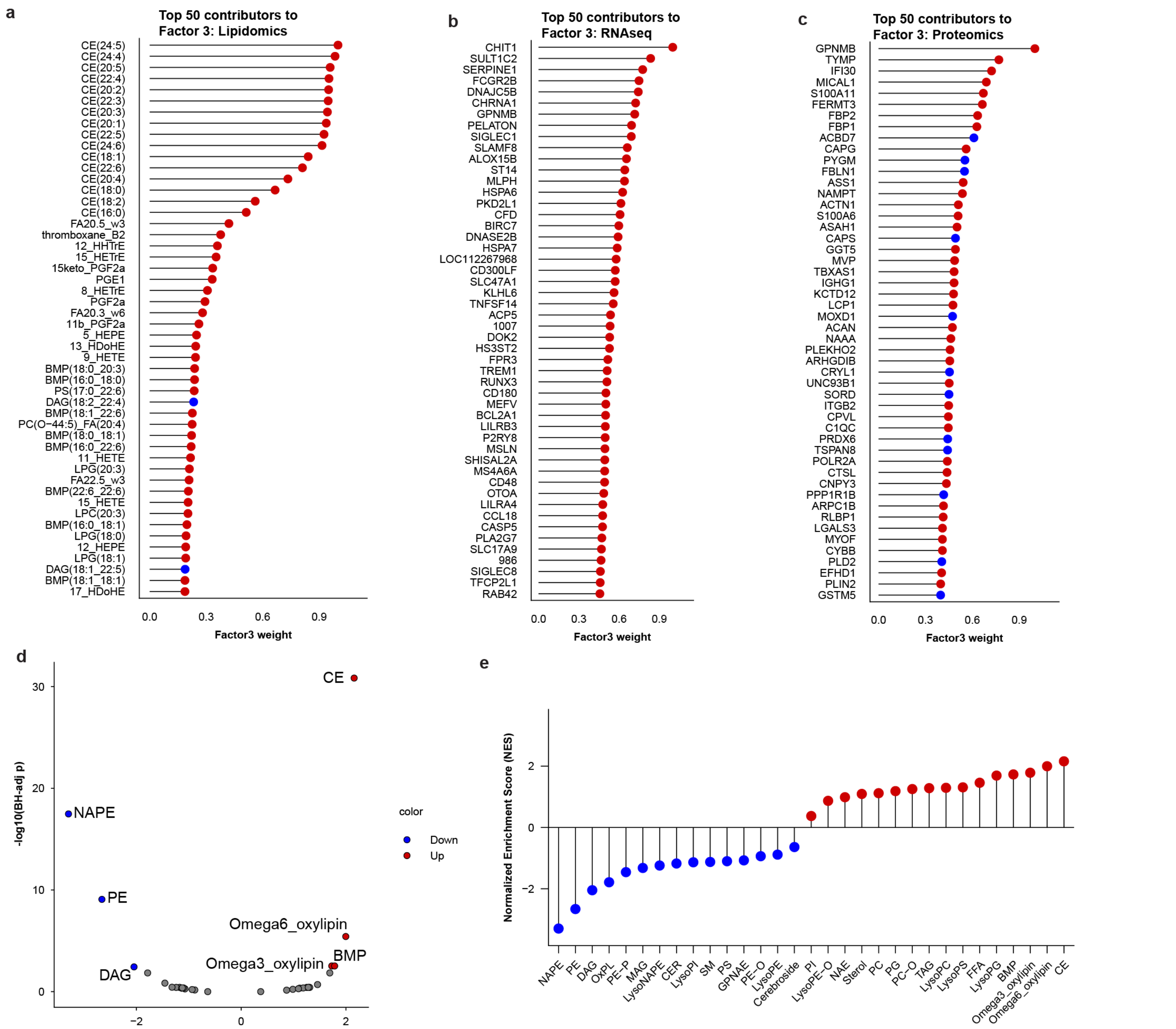


**Figure S15**

**Fig. S19. Top loadings of MOFA factor 3. a-c,** Top 50 lipid (**a**), gene (**b**) or protein (**c**) relative loadings of MOFA factor 3. Data is expressed as the relative loading while the color indicates directionality (red: positively associated with factor 3, blue: negatively associated with factor 3). **d-e,** enrichment of lipid classes enriched in factor 3 using GSEA with the lipid class as custom "genesets".

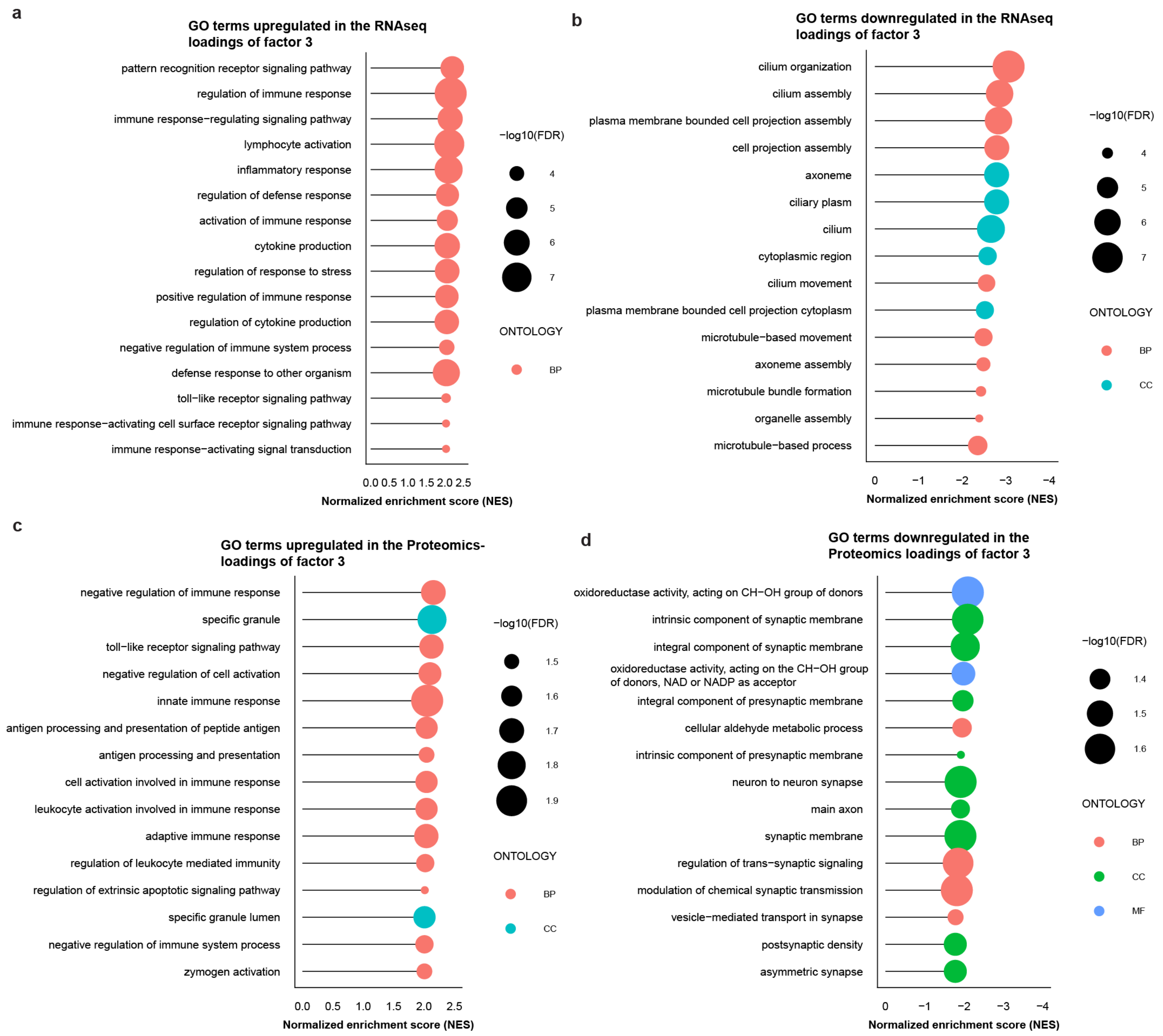


**Fig. S20. Pathways associated with factor 3.**

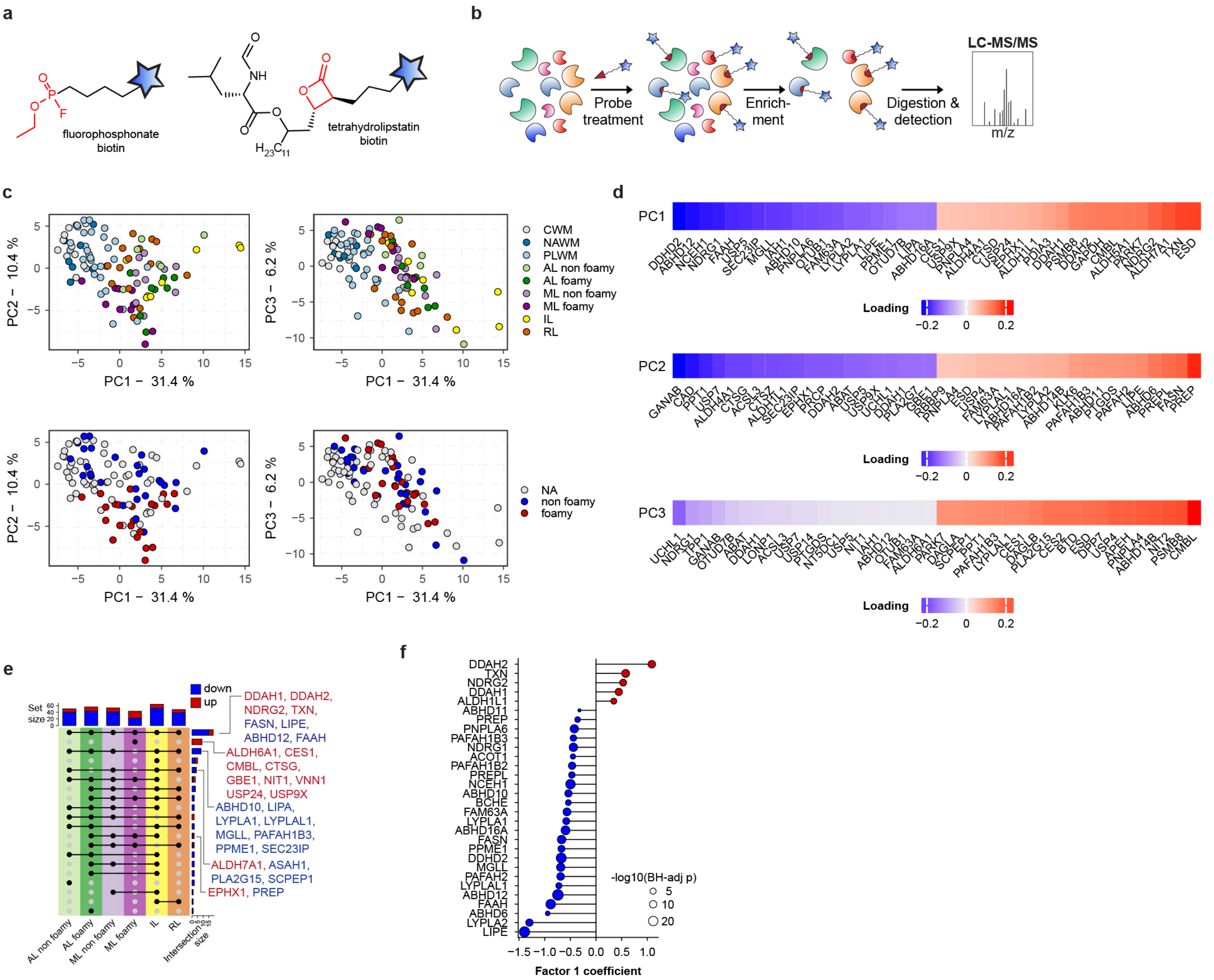

**Fig. S21. Activity-based protein profiling of MS lesions. a,** Probes used for ABPP in this study. **b,** schematic workflow of the ABPP method. Lysates are incubated with the probes from **(a),** and subsequently targeted proteins are enriched from the mixture and analysed by mass spectrometry. **c,** PCA analysis of the ABPP data coloured by lesion type or morphology. **d,** Top 20 positive and negative loadings from the ABPP PCA in **(c). e,** upset plot show overlap of differentially active enzymes in the different lesion type compared to NAWM. **f,** association of enzyme activities with MOFA factor 1. A linear model of the seven MOFA factors was fitted to the ABPP data, identifying enzymes significantly associating to factor 1.

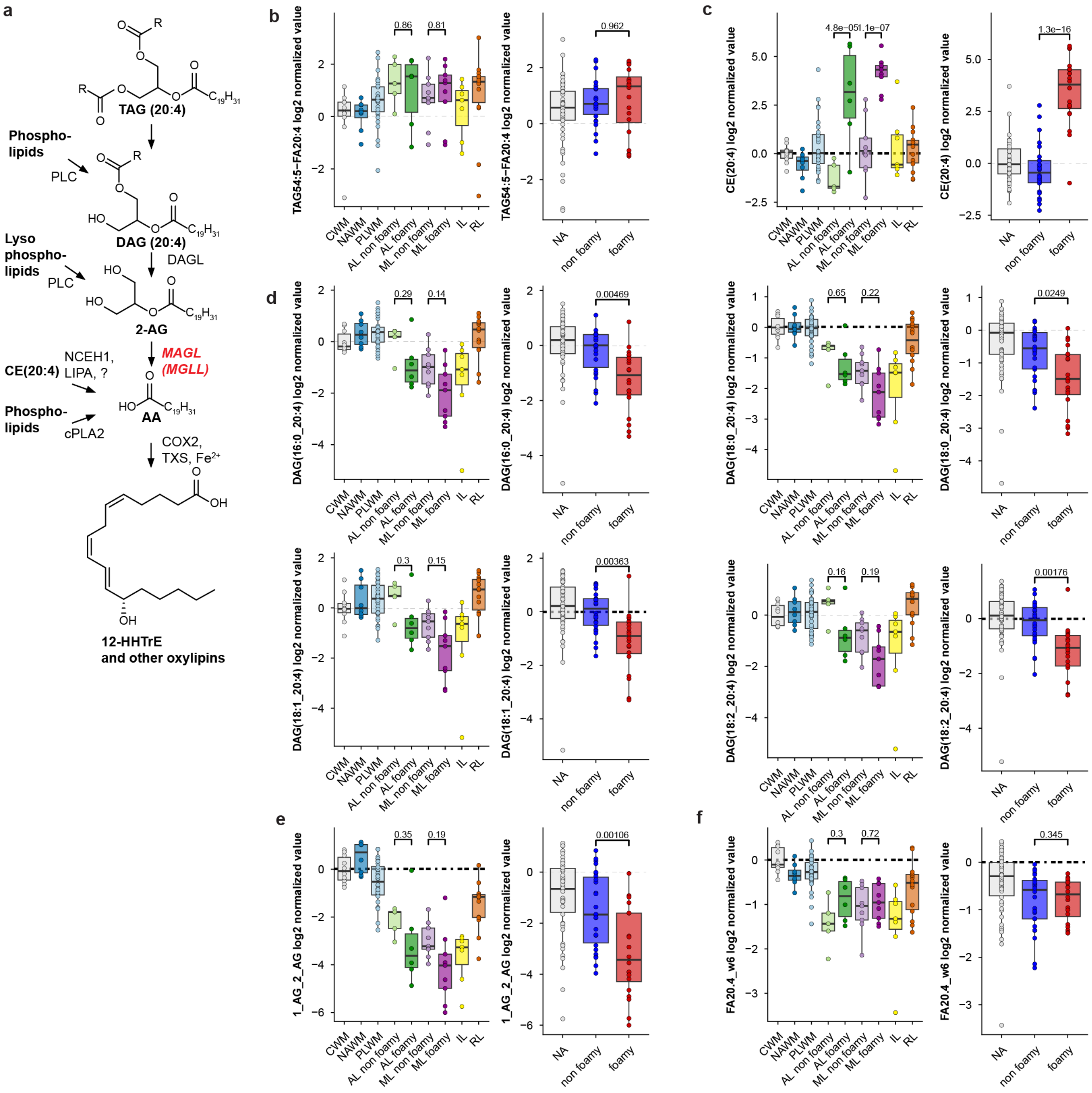


**Fig. S22. Metabolic pathways for the production of free arachidonic acid. a,** the pathway starting from AA-containing TAG, which is hydrolysed to DAG, MAG and subsequently AA. MAGL plays a rate-limiting role in the supply of AA for oxylipin production by controlling the conversion of 2-AG to AA. **b-f,** boxplots indicating the levels of lipids indicated in **a**. statistics are from limma with BH-correction for multiple testing. TAG levels are increased in all lesion types, but AA-containing DAG and 2-AG are decreased in lesions with foamy microglia compared to non-foamy, while oxylipin production was upregulated suggesting an increased flux to arachidonic acid.

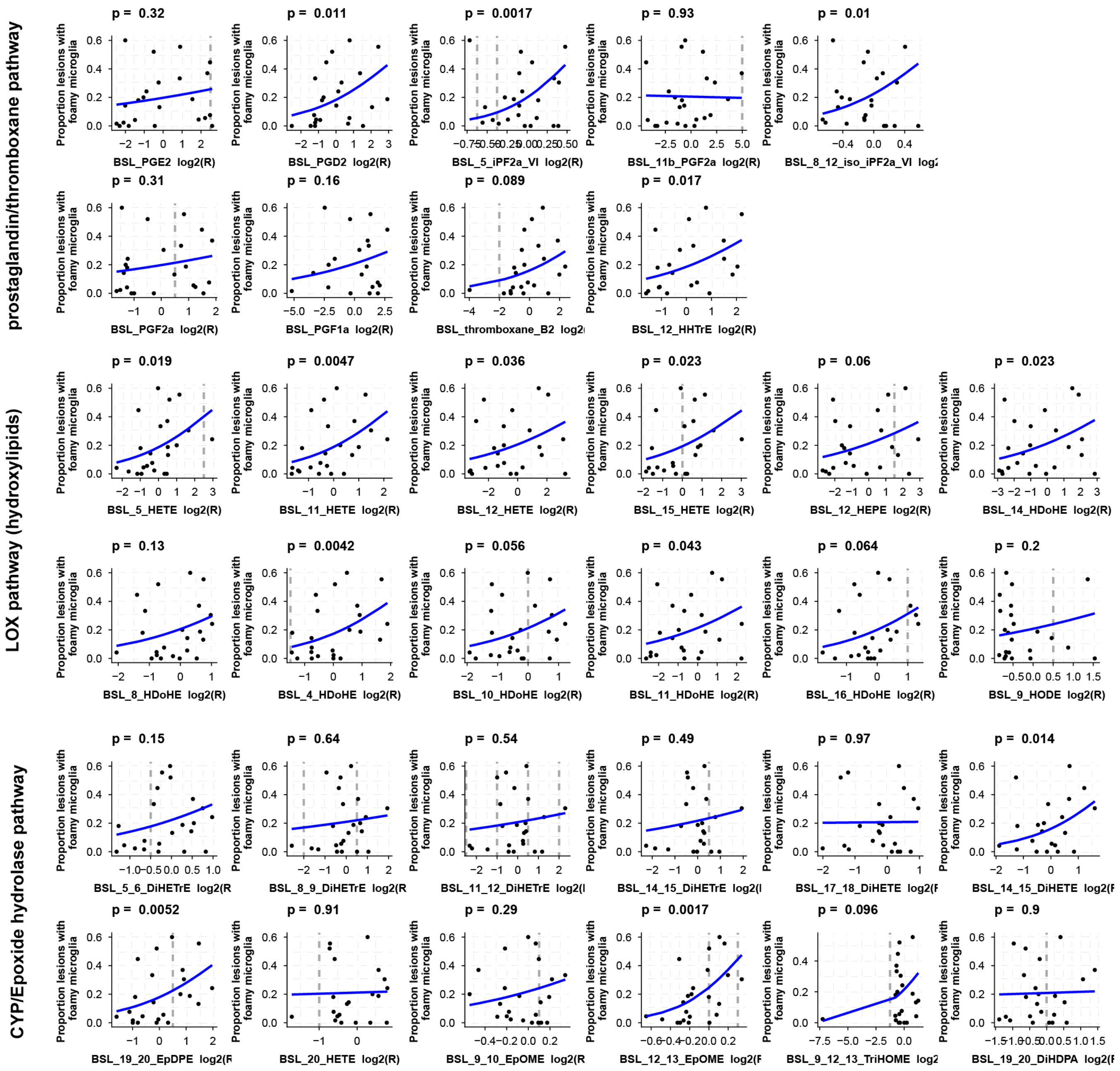


**Fig. S23.** A generalized linear model showing the association between CSF oxylipins to the proportion of foamy lesions, p-values are from a likelihood-ratio test. P-values were not corrected for multiple testing.
